# Physiological α-synuclein S129 phosphorylation mediates postsynaptic and nuclear interactions in the human brain

**DOI:** 10.1101/2025.04.22.649861

**Authors:** Solji G. Choi, Jayda B. Duvernay, Atousa Bahrami, Tyler Tittle, Sepehr Sani, Jeffrey H. Kordower, Scott Muller, Bryan A. Killinger

## Abstract

Phosphorylation of alpha-synuclein (αsyn) at serine 129 (PS129) marks aggregates in synucleinopathies but also occurs physiologically, potentially signaling protein interactions during neuronal activity. Technical barriers, including postmortem dephosphorylation, have hindered the study of physiological PS129 in the human brain. Using biotinylation by antibody recognition (BAR) on surgically resected temporal lobectomy tissues (without post-mortem interval), we mapped physiological PS129 and total αsyn interactomes. BAR identified 1,095 interactions with 513 αsyn-specific, 524 shared, and 58 PS129-specific, mostly associated with vesicles at presynaptic nerve terminals. PS129-specific interactions were uniquely associated with postsynaptic density proteins SHANK1/3, DLGAP1-4, DLGAP1-3, and DLG2-4, as well as nuclear-associated proteins HUWE1, HNRNPM, RBM14, ITCH, OGT, PHF24, and PPP2R5E. Fluorescent staining confirmed physiological PS129 proximal to dendrites and within the nucleus. Confirmation in healthy cynomolgus macaques (62% αsyn and 41% PS129 overlap) demonstrated that the interactomes were physiological rather than disease- or aggregate-associated. We conclude that physiological PS129 plays a unique and underappreciated role in postsynaptic neurons extending from the postsynaptic active zone to the nucleus. These interactomes benchmark normal αsyn biology, illuminating the transition to synucleinopathy pathology.

**Significance Statement:** Disease-associated αsyn phosphorylation (PS129) was recently identified in healthy mammalian brain and may signal αsyn-protein interactions during neuronal activity. Here, we surmounted technical hurdles and characterized αsyn and PS129 interactomes directly in the human brain. Results showed a unique significance for PS129 in post-synaptic active zones and nuclear compartments, which was confirmed in healthy non-human primates. These αsyn interactomes will be a valuable reference for understanding synucleinopathy mechanisms in the context of normal αsyn biology.

## Introduction

Alpha-synuclein (αsyn) is an intrinsically disordered protein that misfolds and accumulates in neurodegenerative diseases referred to as synucleinopathies(1). Although its precise biological role is still not fully understood, αsyn is concentrated at presynaptic nerve terminals, where it interacts with the SNARE complex and may play a role in vesicle endo/exocytosis(2–10). αSyn is found at lower concentrations in postsynaptic dendrites(11) and nucleus(12), significant sites for synucleinopathy disease mechanisms(13–20). Although elucidating αsyn’s physiological role in synaptic communication is essential for understanding the disease mechanism, relatively few studies have examined αsyn in a non-disease context.

αSyn can be phosphorylated at serine 129 (PS129), which is a specific marker of αsyn aggregates in post-mortem synucleinopathy brain (21, 22). However, PS129 also occurs in the healthy mammalian brain, results from neuronal activation, and may play a role in signaling protein-protein interactions (21, 23–25). Specifically, PS129 increases interactions with VAMP2 and the cytoskeleton(24, 25) and may be related to folding and activity at vesicle membranes(23, 26). PS129 is predominantly found at presynaptic terminals but is also found in dendrites and the nucleus(21, 27, 28). PS129 may play an important role at presynaptic terminals, fine-tuning neurotransmitter release(23, 29) but is also distributed throughout neurons within several subcellular compartments, raising important questions about alternative roles for PS129. Identifying physiological-PS129-specific interactions will help understand their role in biology and the transition to disease.

Technical limitations have prevented physiological-PS129 characterization in the human brain(22, 30). Physiological PS129 is rapidly dephosphorylated following death (22, 31), making post-mortem PS129 measurements unreliable. Theoretically, brain tissues lacking a post-mortem interval (PMI) would be required to study physiological PS129 in its native context.

Post-mortem brain specimens lacking PMI are impractical, and thus surgically resected brain specimens are the only option to study physiological PS129 in the human brain. Temporal lobectomies are performed in rare intractable cases of epilepsy where a small portion of the temporal cortex, hippocampus, and amygdala are surgically resected(32). In the mammalian brain, PS129 is abundant in the hippocampus (21, 24), making temporal lobectomy tissues an optimal source for establishing human PS129.

Biotinylation by antibody recognition (BAR) is an in situ proximity proteomics method used to identify protein interactions within the intact brain(33–35) that has been used to describe alpha-synuclein aggregate interactions (BAR-PS129) and total αsyn interactions (BAR-αSyn) in synucleinopathy brain (34, 35). Physiological PS129 interactions have been described in the olfactory bulb of healthy rodents (21), in primary cells (24), and in postmortem brain tissue lysates (22). αSyn interactomes have been measured in yeast(36–38), primary rat neurons(39), HEK293 cells(40), and post-mortem brain(34). To date, the interactions of physiological PS129 have remained unexplored in the intact human brain in the absence of synucleinopathy.

## Materials and Methods

### Tissues

The Institutional Review Board and the Institutional Animal Care and Use Committee at Rush University Medical Center approved all procedures. Free-floating PD/DLB hippocampal sections were obtained from the Rush Movement Disorders Brain Bank. Formalin-fixed paraffin-embedded (FFPE) hippocampal sections from non-synucleinopathy and PD cases with short postmortem intervals (2-3 hours) were obtained from the Brain and Body Donation Program at Banner Sun Health Research Institute. Surgically resected brain specimens were collected from patients with intractable epilepsy. Details for all human specimens used in these studies are shown in Table 1 (n=5, 1-4 specimens per case). Cynomolgus macaque and C57BL/6J mice (RRID: IMSR_JAX:000664) used in this study are detailed in Table 2.

### Surgical tissue collection and fixation

Upon tissue resection from epileptic patients, samples were either immediately placed in 4% paraformaldehyde (PFA) solution in 0.1 M phosphate-buffered saline (PBS, pH 7.4) or first stored in PBS for a few minutes, then transferred to 4% PFA. The tissues underwent fixation for at least 72 hours, followed by equilibration in 15% and then 30% sucrose solutions. Tissues were then coronally sectioned at 40 μm using a freezing-stage sliding knife microtome (American Optical) (RRID:SCR_018434). Sections were stored in a cryoprotectant solution (30% sucrose, 30% ethylene glycol in PBS) at −20 °C until further processing. Tissues that were too small for cryosectioning or small remnant tissues after cryosectioning were processed for FFPE and sectioned at 4 μm thickness.

### Immunohistochemistry (IHC)

Free-floating sections or FFPE sections (rehydrated through xylenes, graded ethanol, and ultrapure water) were washed three times in dilution media (DM: 50 mM Tris–HCl pH 7.4, 150 mM NaCl, 0.5% Triton-X100) for 10 minutes each wash. Sections then underwent heat-induced antigen retrieval (HIAR) using one of two methods: either sodium citrate buffer (10 mM sodium citrate, 0.05% Tween 20, pH 6.0) for 30 minutes at 90°C, or Tris-EDTA buffer (10 mM Tris base, 1 mM EDTA, 0.05% Tween 20, pH 9.0) for 30 minutes at 100°C. Alternatively, for formic acid antigen retrieval, FFPE sections were incubated in formic acid for 5 minutes, then rinsed in ultrapure water before proceeding to the subsequent steps.

Sections were incubated in peroxidase quenching solution (0.3% hydrogen peroxide, 0.1% sodium azide) containing blocking buffer (3% goat serum, 2% BSA, 0.4% Triton X-100 in DM) for 1 hour at room temperature. Sections were briefly washed in DM and incubated overnight at 4°C with PS129 antibody (Abcam, EP1536Y, RRID:AB_765074). The next day, tissues were rinsed three times in DM for 10 minutes each. Sections were incubated with HRP-conjugated goat anti-rabbit secondary antibody (Thermo Fisher Scientific, Cat# 31462, RRID:AB_228338) at 1:1,000 dilution in DM containing 1% BSA and 1% goat serum for 1 hour at room temperature. Samples were rinsed twice in DM and once in sodium acetate buffer (0.2 M imidazole, 1.0 M sodium acetate buffer, pH 7.2) for 10 minutes each, then developed using a standard nickel-enhanced 3,3′-diaminobenzidine (DAB)-imidazole protocol (dx.doi.org/10.17504/protocols.io.rm7vzx725gx1/v1). Sections were rinsed with sodium acetate buffer and PBS, then mounted on charged slides (Superfrost Plus Microscope Slides, Fisher, Cat# 22-034-979) or gelatin-coated glass slides. After complete drying, sections were counterstained with purified methyl green (Sigma-Aldrich, Cat# 67060), dehydrated, cleared with xylenes (Fisher, Cat# X3P-1GAL), and coverslipped with Cytosol XYL (Epredia, Ref# 8312-4).

### In situ enzymatic pretreatment

Before immunoassays, matching tissues were treated with or without calf intestinal alkaline phosphatase (CIAP) or proteinase K (PK). For the CIAP treatment, tissues were incubated with CIAP (Promega, Cat#: M2825, 20 U/μL) at a 1:333 dilution in CIAP buffer (100 mM NaCl, 10 mM MgCl₂, and 50 mM Tris–HCl at pH 7.9) for 24 hours at 37°C before subsequent assays, following previously established protocol (31). For PK treatment, tissues were incubated with proteinase K (PK, Thermo Scientific, Cat# EO0491, ∼20 mg/mL) diluted 1:1,000 in PBS at 37°C for 10 minutes before proceeding (31).

### Protein extraction and precipitation

For free-floating tissues, proteins were extracted using crosslink reversal buffer (5% SDS, 0.5 M EDTA pH 8.0, 0.5 M Tris-base, 0.150 M NaCl) as described previously (35) (see protocols.io https://dx.doi.org/10.17504/protocols.io.36wgq3xyylk5/v1). Following extraction, 100 μL of the sample was mixed sequentially with 400 μL methanol (Sigma-Aldrich, Cat# 179337), 100 μL chloroform (Sigma-Aldrich, Cat# C2432), and 300 μL ultrapure water. After each addition, samples were vigorously vortexed for approximately 10 seconds. The mixtures were centrifuged at 14,000 X g for 2 minutes, and the top aqueous layer was removed. Subsequently, 400 μL of methanol was added to each sample, which was then vortexed and centrifuged at 14,000 × g for 3 minutes. After removing the methanol, the pellets were air-dried for approximately 10 minutes, then resuspended in a 5% SDS solution. For FFPE sections, tissue was first scraped using a clean razor blade and transferred to 1.5 mL Eppendorf tubes.

Deparaffinization and rehydration were performed using graded ethanol and water (41, 42). After centrifuging samples at 21,100 X g for 20 minutes, protein extraction was performed with 50 μL of crosslink reversal buffer using the same method as for free-floating tissues. Next, 50 μL of 8 M urea solubilization buffer (8 M urea, 5 mM dithiothreitol, 150 mM NaCl, 50 mM Tris-HCl, pH 7.5) was added to the samples, which were then sonicated using a probe sonicator (Misonix, Microson Ultrasonic Cell Disruptor XL). Samples were centrifuged at 21,100 × g for 20 minutes, then the protein was precipitated and resuspended in 5% SDS solution.

### Spot blotting

1 µL of each protein sample was spotted onto a methanol-activated PVDF membrane and dried. The membrane was reactivated with methanol, washed with ultrapure water, and then fixed in 4% PFA for 30 minutes. The blots were subsequently rinsed in water and probed for total protein stain (Licor, Revert 700 Total Protein Stain Kit, Part# 926-11010). Upon imaging the total protein with Odyssey M imager (Li-Cor) (RRID:SCR_025709), blots were destained using Revert 700 destaining solution, followed by rinsing in TBST (20 mM Tris–HCl pH 7.6, 150 mM NaCl, 0.1% Tween-20) and placed in blocking buffer (5% dry milk in TBST) for 1 h at room temperature. The membrane was incubated overnight at 4°C with anti-αsyn antibodies: either BD Biosciences (#610787, RRID:AB_398108) at 1:2,000, Abcam EPR20535 (RRID: AB_2941889) at 1:20,000, or Abcam EP1536Y (PS129) at 1:50,000 in blocking buffer. The next day, blots were washed and incubated with secondary antibodies (anti-mouse HRP conjugate at 1:6,000, RRID:AB_330924, and anti-rabbit HRP conjugate at 1:20,000) diluted in blocking buffer. Following TBST wash, blots were imaged using chemiluminescence substrates (Clarity Western ECL Substrate, Bio-Rad, product# 170–5060 or Super Signal West Atto Ultimate Sensitivity Chemiluminescence Substrate, Thermo Scientific, Ref# A38554) with a Chemidoc Imager (Bio-Rad, RRID:SCR_019037). For quantification of signal intensities, the optimal auto-exposure setting was applied during imaging, and the resulting raw TIFF files were analyzed using ImageJ (version 1.54g, RRID:SCR_003070). The results were graphed, and a Two-Way ANOVA with Tukey post hoc testing was performed in GraphPad Prism 10. Western blotting 10 µg of proteins were separated using 4–12% Bis–Tris gels (Invitrogen, Ref# NW04125BOX) with MES SDS Running Buffer (Invitrogen, Ref# B002) and blotted onto methanol-activated polyvinylidene difluoride (PVDF) membrane (Immobilon-FL Membrane, Merck Millipore Ltd, Cat# IPFL00010). The membrane dried and reactivated with methanol, then fixed in 4% PFA for 30 min, rinsed twice in ultrapure water, stained for total protein, rinsed in water and TBST, and blocked with blocking buffer (5% dry milk in TBST) for 1 h at room temperature. Blots were probed and imaged as described for spot blotting.

### Quantification of the immunohistochemistry

PS129 was quantified for dark black pixels using the Ilastik (RRID:SCR_015246) pixel classification workflow. All images were imported into Ilastik and trained to distinguish signal from the background. Then, batch analysis with the established parameters was applied to all images simultaneously without user interaction (43). The pixel classification-processed images were saved and subsequently analyzed in ImageJ for quantification. A macro was recorded using the following steps (44): upon opening an image, changing the LUT to Glasbey, applying auto-threshold with MaxEntropy, and measuring the area containing signals. This recorded macro was applied identically to all images, and the resulting area measurements were exported to Excel and graphed in Prism to represent the percentage of signal area.

### BAR

Proximity labeling experiments were performed using primary antibodies against PS129 (BAR-PSER129, Abcam EP1536Y), total αsyn (BAR-αSyn, Abcam EPR20535), and a primary antibody omission negative control (BAR-NEG), as previously described (35). The biotinyl tyramide amplification reaction was conducted in 100 mL of borate buffer (0.05 M sodium borate, pH 8.5) containing 14.7 μL of biotinylated tyramide stock solution (12.5 mg/mL in DMSO; Sigma-Aldrich, SML2135) and 10 μL of 30% hydrogen peroxide (Sigma-Aldrich, H1009) for 30 minutes at room temperature. Following BAR labeling, floating sections were washed with PBS and solubilized in crosslink reversal buffer. Tissues were heated to 98°C for 30 minutes and centrifuged. The resulting supernatant was diluted in TBST buffer (150 mM NaCl, 1% Triton X-100, 50 mM Tris-HCl, pH 7.6), and biotinylated proteins were captured using 40 μL of streptavidin-coated magnetic beads (Thermo Fisher Scientific, Cat# PI88817) by incubation for 1 hour at room temperature with gentle rotation. Beads were isolated using a magnetic stand (Millipore, Magna GrIP Rack, 20-400) and washed two times with wash buffer (150 mM NaCl, 50 mM Tris-HCl pH 7.6, 1.0 mM EDTA pH 8.0, 0.1% SDS, 1.0% Triton X-100) for 30 minutes each, followed by overnight incubation at 4°C for 16 hours. Captured proteins were eluted by heating beads to 98°C for 10 minutes in 50 μL of sample buffer containing 1× LDS sample buffer (Invitrogen, B0008) and 5 mM reducing agent (Roche, 1,4-Dithiothreitol, 10708984001). A total of 40 μL of each sample was loaded onto 4-12% Bis-Tris gels (Fisher Scientific, NW04127BOX) and electrophoresed briefly at 200 V (∼2 minutes) until the samples entered the gel. Gels were placed in fixation buffer (50% ethanol, 10% acetic acid in ultrapure water) for 1 hour until complete gel shrinkage occurred, then rehydrated with ultrapure water and stained with Coomassie blue (Invitrogen, LC6060). The entire lane containing captured proteins was excised and submitted for liquid chromatography-tandem mass spectrometry (LC-MS/MS) analysis. For LC-MS/MS, gel pieces containing embedded proteins were digested with trypsin according to optimized protocols (21).

### LC-MS/MS analysis of BAR-enriched proteins

Tryptic peptide mixtures were analyzed on an Orbitrap Exploris 240 mass spectrometer. Raw MS data from surgical samples were processed using MaxQuant, with searches against the human proteome database (UP000005640) restricted to reviewed Swiss-Prot canonical entries (20,405 proteins). For cynomolgus macaque (CN), MaxQuant was run with combined human database (UP000005640, 20,504 reviewed proteins) and the Macaca fascicularis database (UP000009130), including 23 reviewed and 17,385 unreviewed Swiss-Prot canonical entries (45–47). CN and humans share 97.9% sequence homology for coding regions(48).

Label-free quantification (LFQ) was performed with the MaxLFQ algorithm, which normalizes MS1 peptide intensities across all LC-MS/MS runs using a sample-similarity network. This method enables accurate relative protein quantification by comparing peptides detected across multiple samples and integrating data from different fractions, thereby reducing the impact of missing values on quantitative accuracy (49). The resulting proteinGroups.txt file from MaxQuant was analyzed using LFQ-Analyst (50). Proteins were prefiltered with a minimum number of values required according to equation x / 2 +1, where x is the number of replicates within a condition. Values were then imputed using the missing value procedure from Perseus (51). Protein-wise linear models combined with empirical Bayes statistics with Benjamini-Hochberg correction (adjusted *p*-value cutoff = 0.05) were used for differential abundance analysis, including single-peptide identifications. Proteins identified in each BAR condition from the full dataset (downloaded from LFQ Analyst; Dataset S1 and Dataset S4) were assigned an importance score by multiplying their abundance (log₂ fold-change) by their significance (–log₁₀ adjusted p value), thereby emphasizing both effect size (|log₂FC|) and statistical confidence (Dataset S2, Dataset S4)(52–55). The Human Protein Atlas (version 24.1), UniProt, NCBI Gene database, and GeneCards (version 5.25) were used to retrieve information about the identified proteins.

### Visualization of the BAR-enriched proteins

R (version 4.4.1) was used for data import, statistical analysis, and visualization with the following packages: EnhancedVolcano (volcano plots), readxl (Excel import), dplyr (data manipulation), ggplot2 and ggrepel (scatter and bar plots with labeled points), patchwork (multi-panel figures), RColorBrewer (color palettes), and wordcloud (Gene Ontology term clouds).

Heatmaps, protein importance rankings (dot plots), and GO term bar plots were generated in GraphPad Prism. Weighted Venn diagrams were created using DeepVenn(https://doi.org/10.48550/arXiv.2210.04597). To annotate a display protein-protein interaction networks we used Cytoscape_v3.10.3, stringApp v2.2.0, CentiScaPe v2.2, AutoAnnotate v1.5.2, EnrichmentMap v3.5.0, and clusterMaker2 v2.3.4.(56, 57).

### Multiplex fluorescent labeling

Free-floating tissues were rinsed in DM, then incubated in sodium citrate HIAR at 90 °C for 30 minutes. After cooling, tissue peroxidases were quenched for 1 hour at room temperature, as performed for IHC, and then incubated overnight with PS129 primary antibody (Abcam, EP1536Y, 1:50,000) in blocking buffer. The next day, tissues were washed in DM and incubated with goat anti-rabbit HRP secondary antibody (ThermoFisher Scientific, Cat# G-21040, RRID: AB_2536527, 1:1,000 dilution) for 90 minutes. They were then developed in a tyramide fluorophore solution (borate buffer, 0.003% hydrogen peroxide, 5 µM CF568; Fisher Scientific, Cat# 50-196-4912) for 30 minutes. From this step onward, tissues were protected from light. After two DM washes (5 minutes each), primary antibody was eluted by sodium citrate HIAR. Tissues were blocked and incubated overnight at 4 °C with MAP2 (Abcam, ab302487, 1:10,000 dilution). Following three DM rinses, tissues were incubated with a horse anti-goat secondary antibody (Vector Laboratories, Cat# PI-9500, RRID: AB_2336124) for 90 minutes, then developed with a CF488 tyramide fluorophore solution (5 µM; Fisher Scientific, Cat# NC1474216) for 30 minutes. Tissues were washed three times in DM (5 minutes each), stained with DAPI (Invitrogen, Cat# D1306, 1:2,000) for 30 minutes, washed three times in PBS (5 minutes each), mounted on charged slides, and cover-slipped with anti-fade mounting medium (Abcam, ab104135).

## Results

### PMI and tissue processing are critical factors for detecting physiological PS129

Physiological PS129 occurs in healthy mammalian brain but has not been unambiguously identified in non-synucleinopathy human brain, potentially due to post-mortem dephosphorylation (22, 31). Our previous mouse brain studies demonstrated that rapid brain fixation was critical for detecting physiological PS129, as it prevents post-mortem dephosphorylation of αsyn (31). To date, we’ve observed evidence of human physiological-PS129 in the olfactory bulb of a single low-PMI case(21). To determine whether physiological PS129 occurs in the human brain, we obtained both non-synucleinopathy (i.e., control) and PD formalin-fixed paraffin-embedded (FFPE) tissues containing hippocampus with markedly short PMIs (∼2-3 hours) from the Banner Sun Health Research Institute (see Table 1 for specimen details). We performed IHC on these FFPE sections using sodium citrate buffer (pH 6.0) and heat-induced antigen retrieval (HIAR), an effective method for detecting physiological PS129 in the mouse brain (31). Unexpectedly, PS129 reactivity was completely absent in the non-synucleinopathy hippocampus. PS129 reactivity was observed in PD hippocampus even following CIAP pretreatment, indicating aggregated PS129 (Fig. 1A, Fig. S1) (31). To assess whether the physiological PS129 epitope was masked, we applied more stringent antigen retrieval methods (Tris-EDTA and formic acid); however, these approaches also failed to detect PS129 in non-synucleinopathy hippocampal specimens (Fig. 1A. Fig. S1).

**Figure 1.**
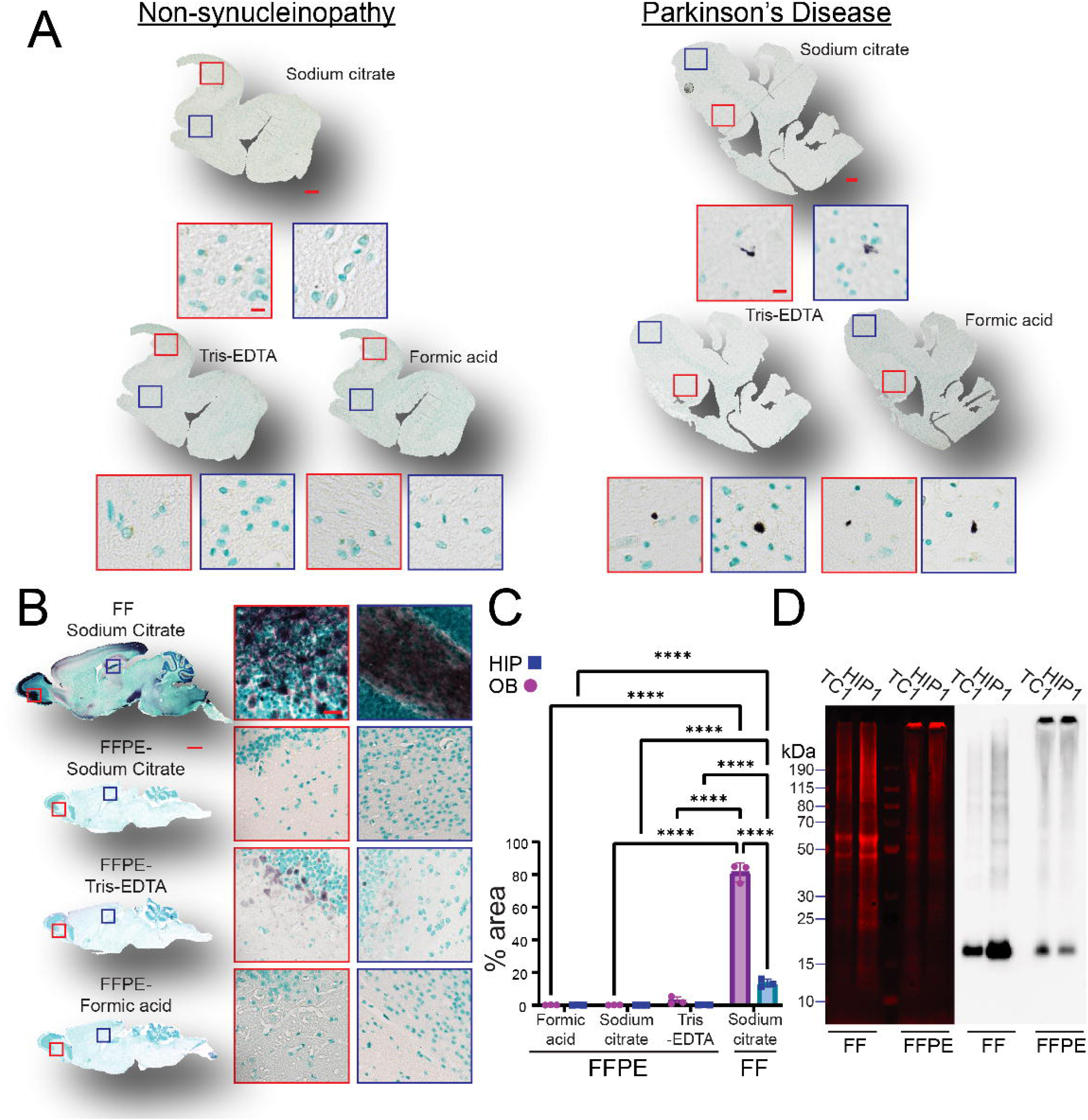
Physiological PS129 is not reliably detectable in FFPE sections. (A) IHC for PS129 in hippocampus-containing temporal cortex from control (non-synucleinopathy) and PD cases, pretreated with sodium citrate, Tris-EDTA, or formic acid for antigen retrieval. Enlarged 20× images correspond to red- and blue-boxed regions in whole-tissue scans. Nickel-enhanced DAB (black); methyl green counterstain. Scale bars = 2 mm and 10 μm. (B) Three mice perfusion-fixed with 4% PFA; one brain half processed free-floating, the other as FFPE. Three antigen retrieval methods applied before PS129 IHC. Representative whole-tissue and 20× images shown for olfactory bulb (OB) and hippocampus (HIP). Scale bars = 1 mm and 25 μm. (C) OB and HIP quantified for % PS129-positive area in 20× images using Ilastik and ImageJ (mean ± SD; Šidák’s multiple comparisons test, ****adj. p < 0.0001). (D) 10 μg protein lysates from free-floating and FFPE sections (surgical TC1, HIP1) separated on 4–12% Bis-Tris gels, transferred to PVDF, and probed for total protein and PS129.

Despite remarkably short PMIs, PS129 was undetectable in post-mortem FFPE tissues; thus, additional processing factors likely interfered with PS129 detection. To test this possibility, we optimally perfusion-fixed (31) wild-type mice, bisected the brains sagittally, and processed one hemisphere as free-floating and the matching hemisphere as FFPE format. Results were striking; free-floating tissues had robust PS129 immunoreactivity, particularly in the olfactory bulb, hippocampus, substantia nigra, and select cortical regions (Fig. 1B). In stark contrast, PS129 reactivity was nearly absent in FFPE sections (Fig. 1B). Among FFPE samples, only those subjected to Tris-EDTA HIAR showed faint PS129 reactivity, restricted to regions typically enriched in PS129, such as the olfactory bulb and hippocampus. Quantification confirmed a significant difference in reactivity between the two formats (F(3,12)=685.3, PL<L0.0001, two-way ANOVA) (Fig. 1C). PS129 reactivity for free-floating was orders of magnitude above FFPE, and not attributable to differences in section thickness (40-micron vs. 4-micron, respectively) but instead suggested epitope masking for FFPE. Blots of extracted proteins from floating sections revealed monomeric PS129 (surgical temporal cortex 1: TC1 and surgical hippocampus 1: HIP1; Fig. 1D), but in FFPE sections, a mixture of monomeric and high molecular weight PS129 was detected. High molecular weight PS129 likely reflects intact formalin crosslinks or insoluble material, typical of proteins extracted from FFPE tissues (42, 58–61). Together, these results demonstrate detection of physiological PS129 in human brains requires rapid fixation and, to some extent, a free-floating format. Unlike animal models, human brain tissue is subject to unavoidable PMIs and slow PFA penetration(62), creating substantial physiological-PS129 detection challenges for post-mortem tissues.

### Physiological, non-aggregated, PS129 is abundant in the human brain

Given requirements (Fig. 1)(31) to detect physiological PS129 in the human brain, we collected surgically resected brain tissues from anterior temporal lobectomy from intractable epilepsy patients. Resected tissues typically included portions of the inferior and medial temporal gyri, covering the hippocampus and the amygdala anterior to the hippocampus.

Following resection, tissues were either rapidly fixed in 4% PFA or briefly immersed in saline before fixation (i.e., all tissues except for TC5 had fixation started in PFA within 3 min. Details of fixation for all cases are provided in Table 1). These procedures provided the shortest possible PMI for human brain specimens in a brain region where physiological PS129 is likely to be abundant (21, 31).

In these specimens, IHC revealed strong PS129 positivity in both the resected hippocampus (Fig. 2A, Fig. S2A) and temporal cortex (Fig. 2B, Fig. S2A) only in free-floating sections (Fig. S3), with PS129 predominantly throughout the gray matter. There were intensely labeled PS129-positive perikarya; with some cells, PS129 was more prominent in nuclei than cytoplasm, and several neurons exhibited distinct PS129-positive projections extending from their cell bodies. Diffuse punctate PS129-positive structures were across PS129-reactive regions, consistent with localization at presynaptic terminals (63). CIAP and PK pretreatment abolished PS129 staining in the surgical hippocampus and temporal cortex (Fig. 2A and 2B), confirming that phosphorylated αsyn was non-aggregated(31). In stark contrast, in the PD brain, CIAP pretreatment did not abolish PS129 staining (Fig. 2C). In the hemisection shown, the hippocampus (Fig. 2C, red box) and the mid-inferior temporal cortex (Fig 2C, blue box) as well as other cortical regions (e.g., cingulate) showed dense Lewy bodies and Lewy neurites (64).

**Figure 2.**
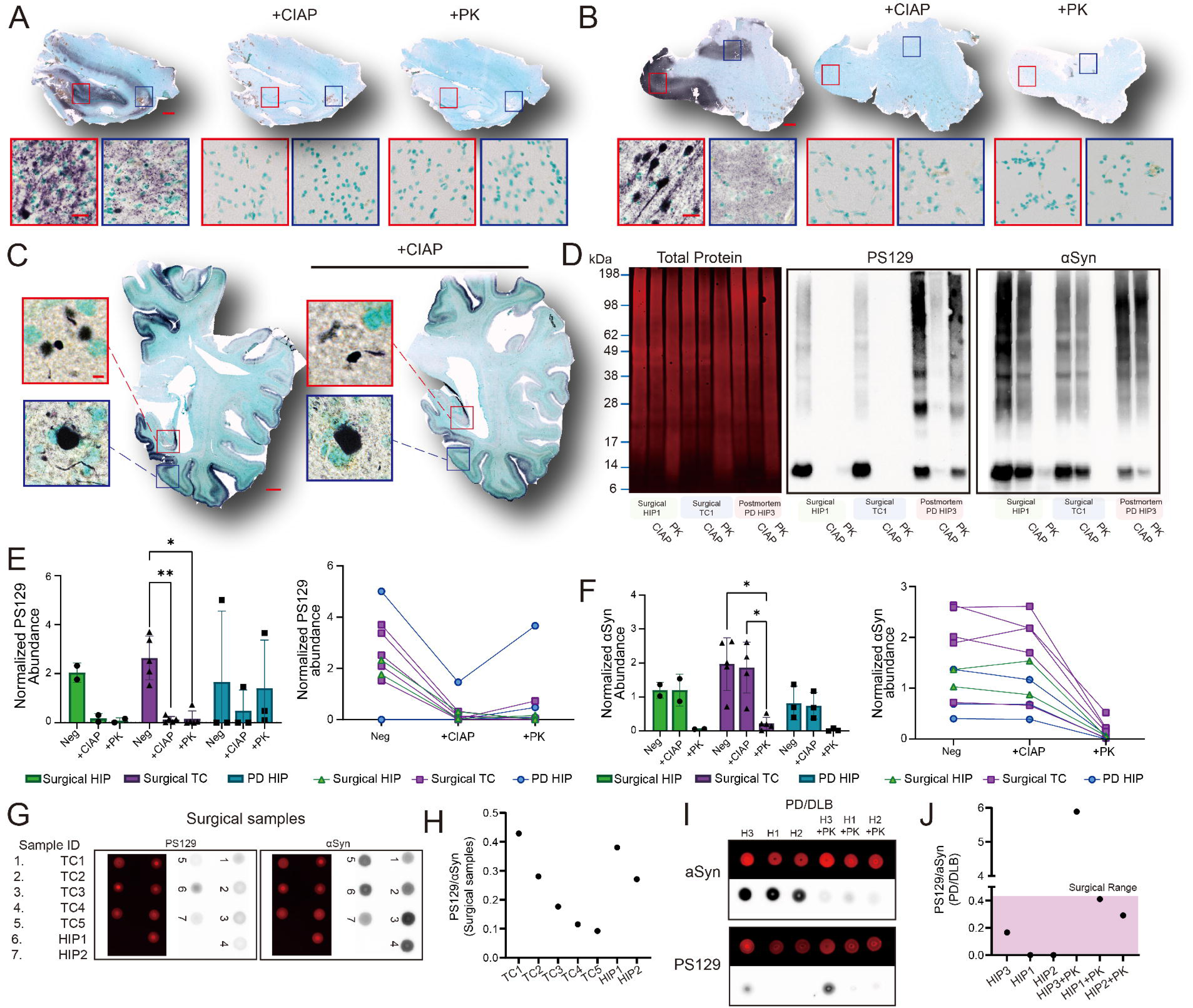
PS129 in surgical samples is a CIAP-sensitive, monomeric form of αsyn. (A), (B). PS129 immunoreactivity is detected in specific regions of surgical samples (Sample IDs: HIP2, TC5; Table 1) and is abolished by CIAP or PK treatment. (C) In contrast, PD hemisections (Sample ID: PD HIP3) show aggregated, CIAP-resistant PS129 predominantly in gray matter. Scale bars: A, B = 1 mm and 12.5 μm; C = 1 cm and 3 μm. (D) Tissue lysates (10 μg protein per sample) were separated on a 4–12% Bis–Tris gel, transferred to PVDF membranes, and stained for total protein (Revert 700, LI-COR), PS129, or αSyn. Representative blots show total protein and chemiluminescent signals for PS129 and αSyn in surgical HIP, TC, and PD HIP samples (Sample IDs: HIP1, TC1, PD HIP3). Negative controls (Neg) denote samples without enzymatic pretreatment; CIAP and PK indicate pretreatment with the respective enzymes. (E) (F). PS129 and αSyn band intensities were normalized to total protein and then to the group mean. Left: normalized protein abundance (mean ± SD; Tukey’s multiple comparisons test, *adj. p < 0.05, **adj. p < 0.01). Right: paired values showing abundance changes within the same samples under different enzymatic conditions. (G–J) One microliter of surgical or PD lysate was spotted onto methanol-activated PVDF membranes and probed for total protein, PS129, or αSyn. Membranes were imaged simultaneously using optimal auto-exposure settings (ChemiDoc, Bio-Rad). Signal intensities were quantified using ImageJ, and PS129:αSyn ratios are shown as scatter plots.

We ran western blot to ensure PS129 specificity, and in surgical lysates, PS129 was mostly monomeric (∼14 kDa) with some high-molecular-weight species (>28 kDa). In contrast, PD hippocampal lysates PS129 ranged from 14 kDa to over 198 kDa, with prominent higher-molecular-weight species (Fig. 2D and Fig. S4A). In surgical samples, CIAP or PK treatment nearly completely abolished PS129 reactivity (Fig. 2E; Tukey’s multiple comparisons: NC vs. +CIAP, mean difference = 2.514, 95% CI [1.259 – 3.769], P = 0.0045; NC vs. +PK, mean difference = 2.481, 95% CI [0.815 – 4.148], P = 0.0133 for five TC samples). In contrast, PS129 reactivity in PD samples persisted or was even enhanced after PK treatment, consistent with our IHC observations (Fig. 2C and Fig. S2B) and previous reports (65, 66). αSyn levels (Fig. 2F) remained unchanged following CIAP treatment in both surgical and PD/DLB cases, while PK treatment rendered αsyn undetectable using the widely used BD SYN1 antibody (RRID: AB_398107) (67) (Fig. 2F; Tukey’s multiple comparisons: NC vs. +PK, mean difference = 1.749, 95% CI [0.5907 – 2.907], P = 0.0126; +CIAP vs. +PK, mean difference = 1.647, 95% CI [0.6129 – 2.680], P = 0.0104, n=5). BDSYN1 recognizes an epitope spanning amino acids 91–99; thus, the epitope may be obscured (e.g., hidden within aggregate conformation) or cleaved in the extractable αsyn (68). Quantification of PS129 and αsyn in seven surgical samples and three PD/DLB hippocampal specimens (Fig. 2E–F) confirmed non-aggregated αsyn in surgical samples and aggregates in PD/DLB specimens.

We then estimated PS129 content in the surgical specimens by spot blotting. Surgical samples exhibited PS129 levels of approximately 10–40% of total αSyn (Fig. 2G, H), with most PD hippocampal samples in the same range (Fig. 2I, J; Fig. S4B). Interestingly, one PD/DLB specimen showed higher PS129 content than αsyn, suggesting that PK destroyed/masked the epitope for the total αsyn antibody (EPR20535), while preserving the PS129 epitope. Further solubilization of insoluble pellets in 8 M urea buffer (22) revealed that surgical samples lacked insoluble PS129 (Fig. S4C), but PD pellets contained CIAP/PK-resistant PS129.

### Physiological PS129 and unphosphorylated αsyn interactomes

BAR was performed as previously described (35) to capture proteins in proximity—primarily within a ∼50–100 nm radius (69)—to PS129 and αsyn. For each case, 2–4 matching sections were pooled for each BAR capture, depending on tissue size (Supplementary Table 1; average combined wet mass of surgical samples: 17.91 ± 6.18 mg). Following BAR, captured proteins (i.e., bead eluent) were spotted onto PVDF membranes and probed for biotin, PS129, and αsyn (Fig. 3A–D). For BAR-PS129 (Fig. 3A, B), αsyn, PS129, and biotin enrichments ranged from 1.5- to 4.5-fold compared to BAR-Neg, except for TC3. TC3 showed the highest overall enrichment, including PS129 (37.3-fold), and was the only case in which the PS129 signal exceeded that of αsyn (7.1-fold) and biotin (12.7-fold), which may reflect the larger tissue mass (25.7mg; Supplementary Table 1) or higher PS129 abundance suggested by IHC (Fig. S2A). For BAR-αsyn, (Fig. 3C, D), all cases showed strong enrichment of αsyn, PS129, and biotin (30-42-fold). Overall abundance across all samples ranged from 85-fold (αsyn in HIP2) to 294-fold (PS129 in TC3) relative to their respective negative controls. Together, these data demonstrate enrichment for BAR-PS129 and BAR-αSyn.

**Figure 3.**
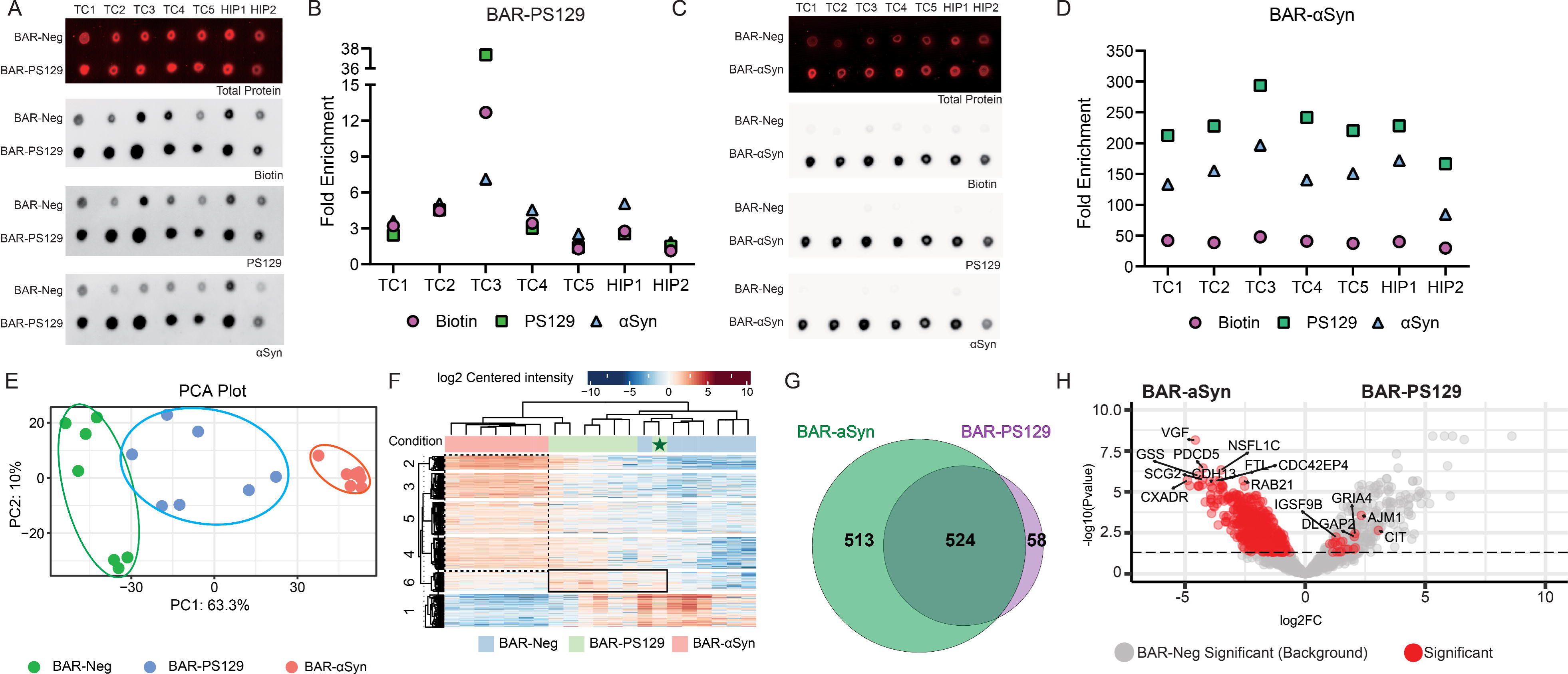
Identification of BAR-αSyn- and BAR-PS129-enriched proteins in surgical samples. All surgical cases were BAR-captured under three conditions: BAR-Neg (primary antibody-omission control, labeled “B-Neg”), BAR-PS129 (“B-PS129”), or BAR-αSyn (“B-αSyn”). (A), (C). A fraction of bead eluate from each sample was spotted onto PVDF membranes and sequentially probed for total protein (Revert Total Protein Stain, LI-COR), biotin (ABC reagent), PS129, and αSyn. (B), (D). BAR enrichment was quantified by normalizing biotin, PS129, or αSyn signal intensities to the averaged relative intensities of negative controls, yielding fold-enrichment values over background. Quantification was performed with ImageJ. Remaining bead eluates were analyzed by LC–MS/MS and quantified using MaxQuant label-free quantification (LFQ). (E) Principal component analysis (PCA) of all samples across the three BAR capture conditions. (F) Heatmap showing unbiased hierarchical clustering of protein abundance across samples. (G) Weighted Venn diagram comparing proteins significantly enriched in BAR-αSyn or BAR-PS129 over background (adj. p < 0.05). A total of 513 and 58 proteins were uniquely enriched in BAR-αSyn and BAR-PS129, respectively, while 524 overlapped. (H) Volcano plot comparing differential protein abundance between BAR-αSyn and BAR-PS129 (adj. p < 0.05). Significant proteins are highlighted in red, with some of the top enriched proteins annotated by name. BAR-Neg significant (background) proteins are shown in gray. Significance thresholds: adj. p < 0.05 and |log₂ fold change| > 1.

BAR-captured proteins were digested with trypsin and analyzed by LC-MS/MS. Principal component analysis (PCA) showed separation between BAR-captures in PC1 and clustering according to BAR-capture (Fig. 3E). PC1 and PC2 explained 73.3% of variance. Hierarchical clustering confirmed similar protein signatures within each BAR-capture group, except for one BAR-PS129 sample (TC5, Fig. 3F, marked with a green star), which clustered with the BAR-Neg group (Fig. 3F). TC5 had the longest delay to fixation (time from tissue resection to immersion in PFA: 10–15 min vs. 0–3 min for others; Table 1), which could reduce protein phosphorylation(22, 31) and subsequent BAR-PS129 enrichment (Fig. 3B), demonstrating the importance of PMI on robust physiological PS129 detection. Protein clustered clearly in the direction of αsyn (Fig. 3F, dotted) and PS129 (Fig. 3F, solid line), and background (Fig. 3F, cluster 1). The complete LFQ-analyst output is provided in Dataset S1 (LFQ full dataset).

A total of 1,095 BAR-αSyn and BAR-PS129 proteins were significantly enriched (i.e., BAR-target vs. BAR-NEG) and compared using a Venn diagram (Fig. 3G). The majority of BAR–enriched proteins overlapped (48%), 513 proteins (47%) were unique to BAR-αSyn, and 58 proteins (5%) were unique to BAR-PS129. BAR-PS129 unique proteins included DLG2–4, DLGAP1–3, SHANK1, GRIA2, GRIA4, IQSEC1, IQSEC2, PCLO, BSN, ERC2, ANKS1B, and RIMS1. Differential abundance between BAR-PS129 and BAR-αSyn (Fig. 3H) revealed 19 proteins significantly more abundant in BAR-PS129 when compared to BAR-αSyn, including GRIA4, AJM1, DLGAP2, IGSF9B, and CIT. For BAR-αSyn, 520 proteins were more abundant than BAR-PS129, including VGF, GSS, PDCD5, NSFL1C, SCG2, FTL, and CDC42EP4.

Background proteins (i.e., significantly enriched in BAR-NEG, Fig. 3H grey) contained abundant mitochondrial biotin-dependent carboxylases such as ACACA, MCCC1, and PCCB, as previously reported(35).

### BAR identifies PS129 and αsyn interaction networks

BAR-αSyn and BAR-PS129 enriched proteins were analyzed cumulatively by STRING to infer protein-protein interaction (PPI) networks. Large highly connected interaction networks were identified with αsyn near the center of both functional and physical interactomes (Figure 4A, SNCA). Functional interaction networks—inferred from shared biological processes (56, 70, 71)—encompassed 18,306 interactions, with ACTB (β-actin) serving as the central hub (i.e., most annotated interactions within the network), with only 2% disconnected nodes (Fig. 4A). In contrast, physical interaction networks—based on direct, experimentally validated protein interactions (56, 70–72)—comprised 5,440 interactions, with UBA52 (ubiquitin A-52 residue ribosomal protein fusion product 1) as the central hub, with 16.2% unconnected nodes. UBA52 contributes to the ubiquitin–proteasome system, autophagy, and is implicated in cellular and animal models of PD (73–76).

**Figure 4.**
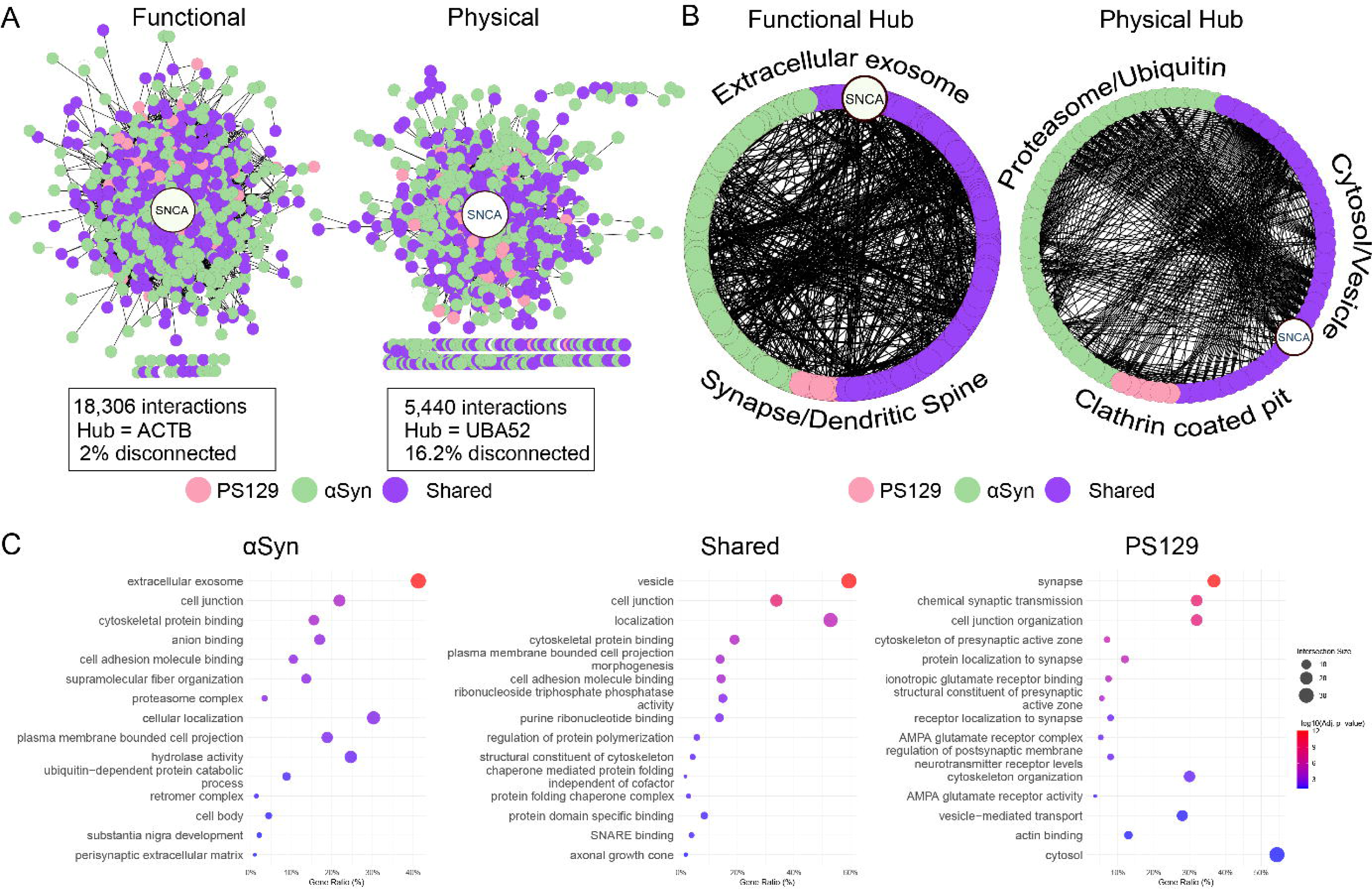
Overall analysis of BAR-αSyn- and BAR-PS129-enriched proteins. (A) Global interaction networks for functional and physical associations. ACTB and UBA52 are identified as the hub proteins for functional and physical interactions, respectively. Nodes represent proteins; edges indicate interactions. (B) Functional and physical interaction maps of central hub protein networks, with edges representing betweenness centrality. Top enrichment terms for αSyn-shared proteins and PS129 proteins are manually annotated within each map. (C) BAR-αSyn and BAR-PS129 were ranked according to their importance scores; an enrichment map of the combined interactome was generated using g:Profiler with ranked queries and visualized in Cytoscape. D. Bar plots of significantly enriched GO terms for ranked BAR-αSyn, BAR-PS129, or shared proteins.

First-degree interactions from hub proteins ACTB and UBA52 were extracted and plotted according to BAR-capture (Fig. 4B). αSyn (SNCA) was a part of these central physical and functional interaction networks. For the functional hub, top GO enrichment terms for shared/asyn interactions involved extracellular exosome, while PS129 interactions involved synapse/dendritic spine. For physical hubs, αsyn proteins enriched for proteasome and ubiquitin, shared proteins cytosol/vesicle, and PS129 interactions involved clathrin-coated pit.

The top 15 enriched GO driver terms for BAR-αSyn, BAR-PS129, and shared proteins (Fig. 4C), were similar to the central network enrichment. BAR-αSyn–enriched terms included extracellular exosome, cell junction, cytoskeletal protein binding, proteasome complex, and hydrolase activity. Shared proteins were enriched for vesicle, cell junction, localization, and SNARE binding. BAR-PS129–enriched terms included synapse, chemical synaptic transmission, cell junction organization, presynaptic active zone cytoskeleton, and several postsynaptic associations.

### Unphosphorylated αsyn associates with the extracellular space

Importance scores (Fig. 5A, Dataset S2) were calculated to reduce dimensionality and rank BAR-enriched proteins(77). For BAR-αSyn, shared and unique proteins distributed across importance score ranks. In contrast, BAR-PS129 highly ranked proteins were mostly shared, with unique proteins appearing at lower ranks. Mean importance scores (Fig. 5B; Dataset S2) were higher for BAR-αSyn (Fig. 5B, Two-way ANOVA F (1, 1615) = 202.3 P<0.0001) as well as differences between unique or shared (F (1, 1615) = 37.05 P<0.0001), suggesting stronger enrichment for shared/overlapping proteins.

**Figure 5.**
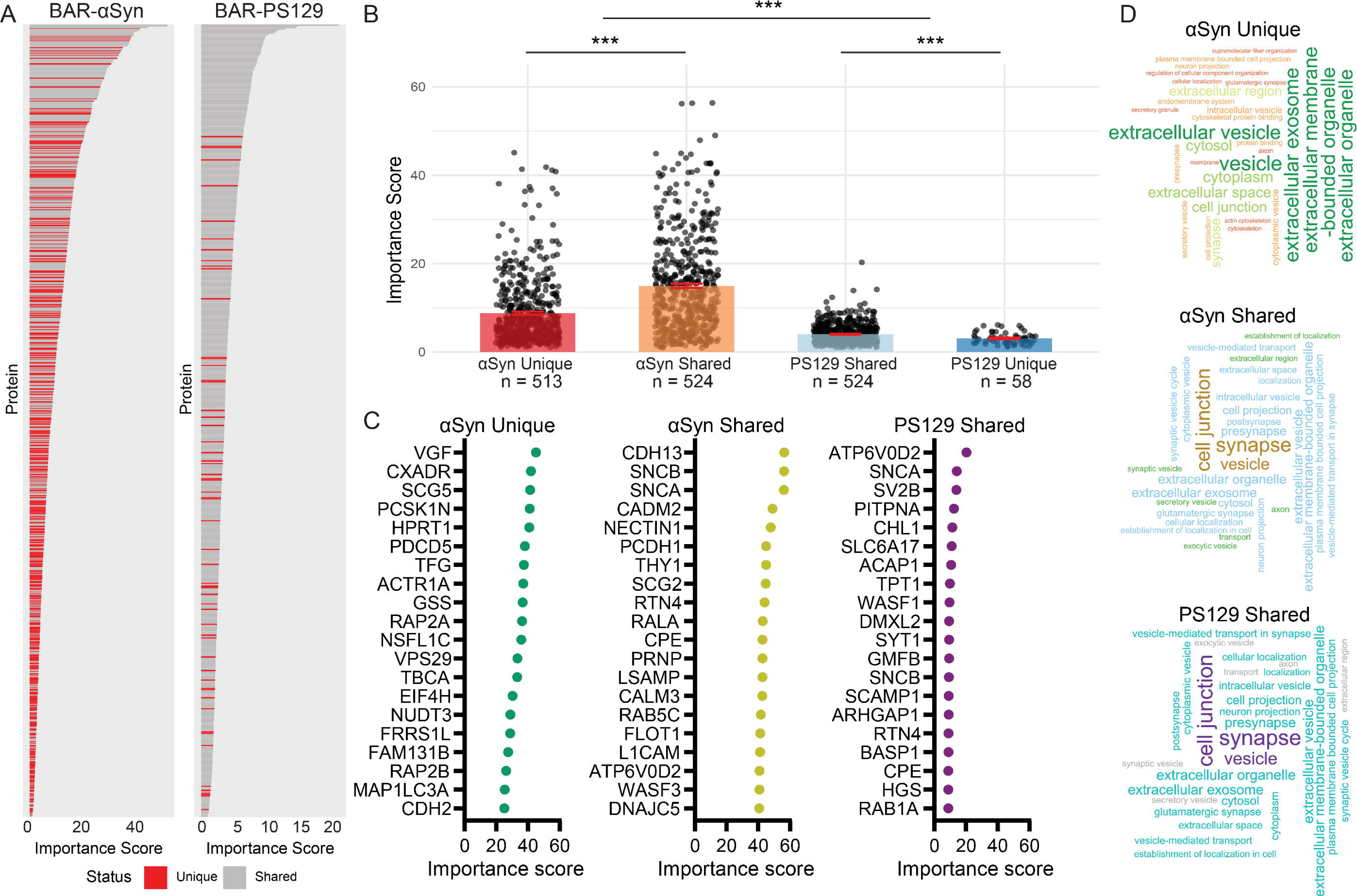
Comparative analysis of unphosphorylated vs. phosphorylated αSyn. (A) Ranked bar plots displaying protein importance scores in the aSyn (left) and PS129 (right) datasets. Proteins are ordered by decreasing importance within each dataset. (B) Mean importance scores for four protein groups: aSyn Unique (n = 513), aSyn Shared (n = 524), PS129 Shared (n = 524), and PS129 Unique (n = 58). Individual data points are jittered for clarity; error bars represent the standard error of the mean (SEM). (C) Dot plots of the top 20 proteins in each group, ranked by importance score. (D) Word clouds of the top 30 enriched GO terms for each protein group, identified through g:Profiler using ranked queries. Font size is proportional to term significance (–log10 adjusted p-value).

In the unphosphorylated αsyn interactome (αsyn-unique), the top 20 proteins included 11 vesicle-associated proteins associated with endosomal and secretory vesicles in the presynaptic compartment or cytoplasm (Fig. 5C). These included VGF (nerve growth factor-inducible; highest-ranked), SCG5, PCSK1N, CXADR, VPS29, NSFL1C, MAP1LC3A, RAP2A, RAP2B, TFG, and ACTR1A. The remaining nine— HPRT1, GSS, EIF4H, NUDT3, FRRS1L, FAM131B, CDH2, TBCA, PDCD5—are intracellular and functioning in metabolism, structural organization, or cell adhesion. Notably, the top protein VGF, has been repeatedly implicated in Parkinson’s disease (78–80).

For αsyn-shared proteins, seven of the top 20 were associated with presynaptic vesicles or vesicle trafficking (Fig. 5C): SNCA, SNCB, ATP6V0D2, CPE, DNAJC5, RALA, and CALM3. The remaining 13 proteins comprised cell adhesion molecules (CDH13, CADM2, NECTIN1, PCDH1, THY1, LSAMP, L1CAM), trafficking, lipid raft, or secretory proteins (FLOT1, PRNP, RAB5C, SCG2), and cytoskeletal regulators (RTN4, WASF3).

For PS129-shared proteins, the top 20 included five that overlap with αsyn shared proteins (ATP6V0D2, CPE, RTN4, SNCA, SNCB). PS129 shared proteins were enriched for presynaptic vesicles or vesicle trafficking, including ATP6V0D2, SLC6A17, SYT1, PITPNA, SCAMP1, SV2B, HGS, RAB1A, CPE, SNCA, and SNCB (11 proteins). The remaining nine proteins—CHL1, WASF1, ARHGAP1, RTN4, BASP1, TPT1, GMFB, DMXL2, and ACAP1—are involved in cytoskeletal remodeling, presynaptic organization, cell projections, and protein homeostasis.

Ranked GO enrichment analysis (i.e., g:Profiler) was performed and top 30 terms used to generate word clouds (Fig. 5D). Among 513 unphosphorylated αsyn-interacting proteins, “vesicle” was the top GO term (284/513 proteins), followed by “extracellular exosome,” “extracellular vesicle,” “extracellular organelle,” and “extracellular membrane-bounded organelle”—all derived from the same 206-protein vesicle subset. Although proteins annotated to these extracellular terms—including Rab GTPases (RAB5A, RAB8A/B, RAB11B, RAB15, RAB35), Arf-family GTPases (ARF4, ARL3, ARL8A/B), and ESCRT components (TSG101, CHMP4B, VPS37C, VTA1)—are predominantly intracellular or presynaptic, they contribute to exosome biogenesis, vesicle secretion, and synaptic release (83–85). Similarly, proteasome subunits, ubiquitin-conjugating enzymes, E3 ligases, and deubiquitinases are intracellular but regulate exosomal cargo sorting and protein quality control (86–88). Although these proteins are synthesized and active intracellularly, they are packaged into extracellular vesicles and detected in the extracellular space (84, 85, 88–90), potentially explaining the strong extracellular enrichment. Although ranking differed between PS129 shared and αsyn shared, enrichment terms were similar.

### PS129-specific proteins associated with the postsynaptic to nuclear axis

For PS129-specific interactions, we gathered proteins exclusively identified in BAR-PS129 (Fig. 3G) or differentially abundant (Fig. 3H) and then ranked by importance score (Fig. 6A). All differentially abundant PS129 proteins were PS129-unique, except FBXO41 (Fig. 6A, highlighted in pink), which was shared with αsyn. 59 PS129-specific proteins were identified, approximately half localized to the postsynaptic density (DLG–SHANK–GRIA–SYNGAP1 network), followed by several proteins associated with the presynaptic active zone and vesicle trafficking (RIMS1, BSN, PCLO, ERC2, DGKQ, DNM3), cytoskeletal proteins (MACF1, SPTBN2, CYFIP1/2, SYNPO), and nuclear/RNA-associated regulatory proteins (HNRNPM, RBM14, HUWE1, OGT, FBXO41, PHF24).The top 15 GO terms for PS129-unique proteins included synapse, postsynaptic density, cell junction, and postsynaptic specialization (Fig. 6B). For differentially abundant proteins, similar patterns were observed, including cell junction, postsynapse, asymmetric synapse, and postsynaptic specialization (Fig. 6B). Enrichment maps comparing PS129-unique and differentially abundant proteins revealed functional interconnectivity centered on synaptic organization (Fig. 6C). The largest clusters shared by both groups included biological quality communication, chemical synaptic transmission, somatodendritic compartment projection, postsynaptic asymmetric density, active zone presynaptic, gated monoatomic ion, and unblocking NMDA calcium, in line with the results shown in Fig. 6B. Notably, although PS129-unique proteins nearly encompass all differentially abundant proteins, terms such as actin filament organization and anatomical morphogenesis differentiation were exclusive to unique proteins, whereas amyloid beta peptide was annotated only for differentially abundant proteins. This distinction likely arises because PS129-unique proteins comprise a larger set and, even for overlapping proteins, the two groups exhibit different ranks for the same proteins.

**Figure 6.**
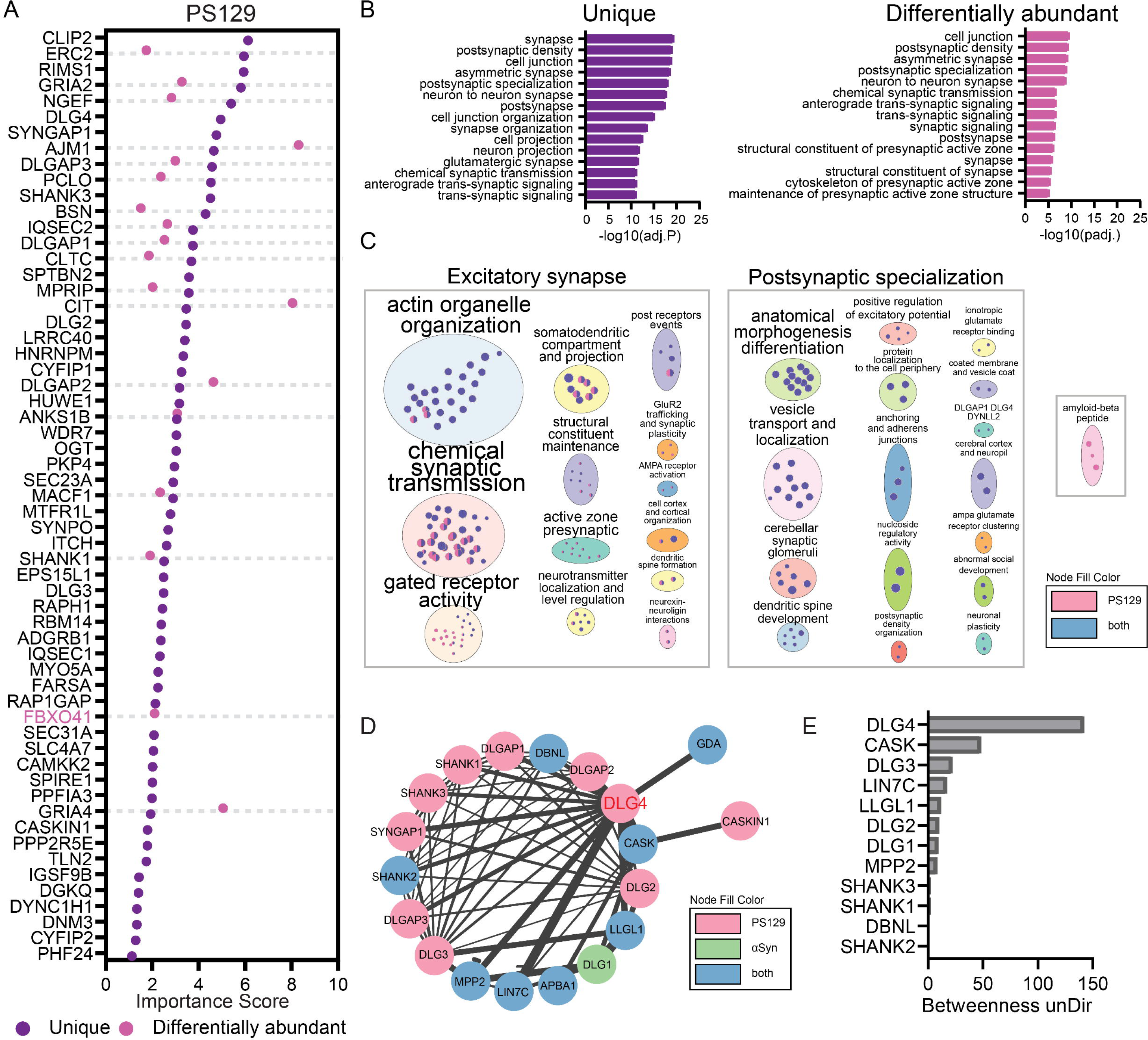
PS129-unique and differentially abundant proteins highlight the PSD as a key distinction from total αSyn. (A) Dot plot of BAR-PS129 proteins. 58 are PS129-unique; 19 are differentially abundant compared to BAR-αSyn (adj. p < 0.05). Ordered by importance; FBXO41 is highlighted in pink, as it was identified only among the differentially abundant proteins in PS129. B. Top 15 enriched GO terms for the 58 PS129-unique or 19 differentially abundant proteins. (C) g:Profiler enrichment analysis of the PS129-unique and PS129-differentially abundant proteins was performed. Results were merged and visualized using Cytoscape Enrichment Map ( FDR cutoff < 0.1, similarity cutoff < 0.25), with node size proportional to the number of genes in each term. For visual clarity, clusters were manually repositioned into each condition. Major overarching themes encompassing the enriched terms were manually annotated atop gray cluster boxes. (D) Cluster identified in the merged BAR-αSyn and BAR-PS129 STRING physical-interaction network (see SF5), with “postsynaptic density” as the top GO:CC term (lowest FDR). (E) Centrality analysis of that cluster: 12 nodes showed measurable undirected betweenness values, with DLG4 (PSD-95) exhibiting the highest.

Merged physical PPI networks of total BAR-αSyn (1,037 proteins) and BAR-PS129 (582 proteins) enriched proteins—followed by Markov Cluster Algorithm (MCL) clustering in Supplementary Figure 5—identified a distinct PSD subnetwork (Fig. 6D) comprising 19 proteins: 10 PS129-unique (DLGAP1–3, SHANK1, SHANK3, SYNGAP1, DLG2–4, and CASKIN1; PSD scaffolds that either are membrane-associated guanylate kinases (MAGUKs) themselves (DLG2–4) or directly or indirectly interact with MAGUK family proteins), 8 shared (SHANK2, DBNL, LIN7C, MPP2, CASK, LLGL1, APBA1, GDA; encompassing MAGUKs and synaptic organizers), and 1 αsyn-unique (DLG1, a MAGUK scaffold). Further analysis of the cluster using betweenness centrality (CentiScaPe) identified DLG4 (PSD-95, a MAGUK family member) as the top hub (score: 142; pS129-unique), critical for PSD organization (91), followed by CASK (calcium/calmodulin-dependent serine protein kinase, a MAGUK) with a score of 48 (Fig. 6E).

### PS129 is proximal to dendrites and within nuclei in the human brain

BAR revealed that PS129 was associated with postsynaptic and nuclear compartments. To validate this, we performed multiplex immunofluorescence on surgical HIP and TC samples, sequentially staining for PS129 and Microtubule-Associated Protein 2 (MAP2), a marker of the somatodendritic domain. Whole-section scan of hippocampus showed robust PS129 signals across all subregions—CA1, CA2, CA3, Dentate Gyrus (DG), and Subiculum (Sub)—with broad overlap in MAP2-positive regions (Fig. 7A, B), as well as distinct PS129 labeling between regions (Fig. 7B, dashed line).

**Figure 7.**
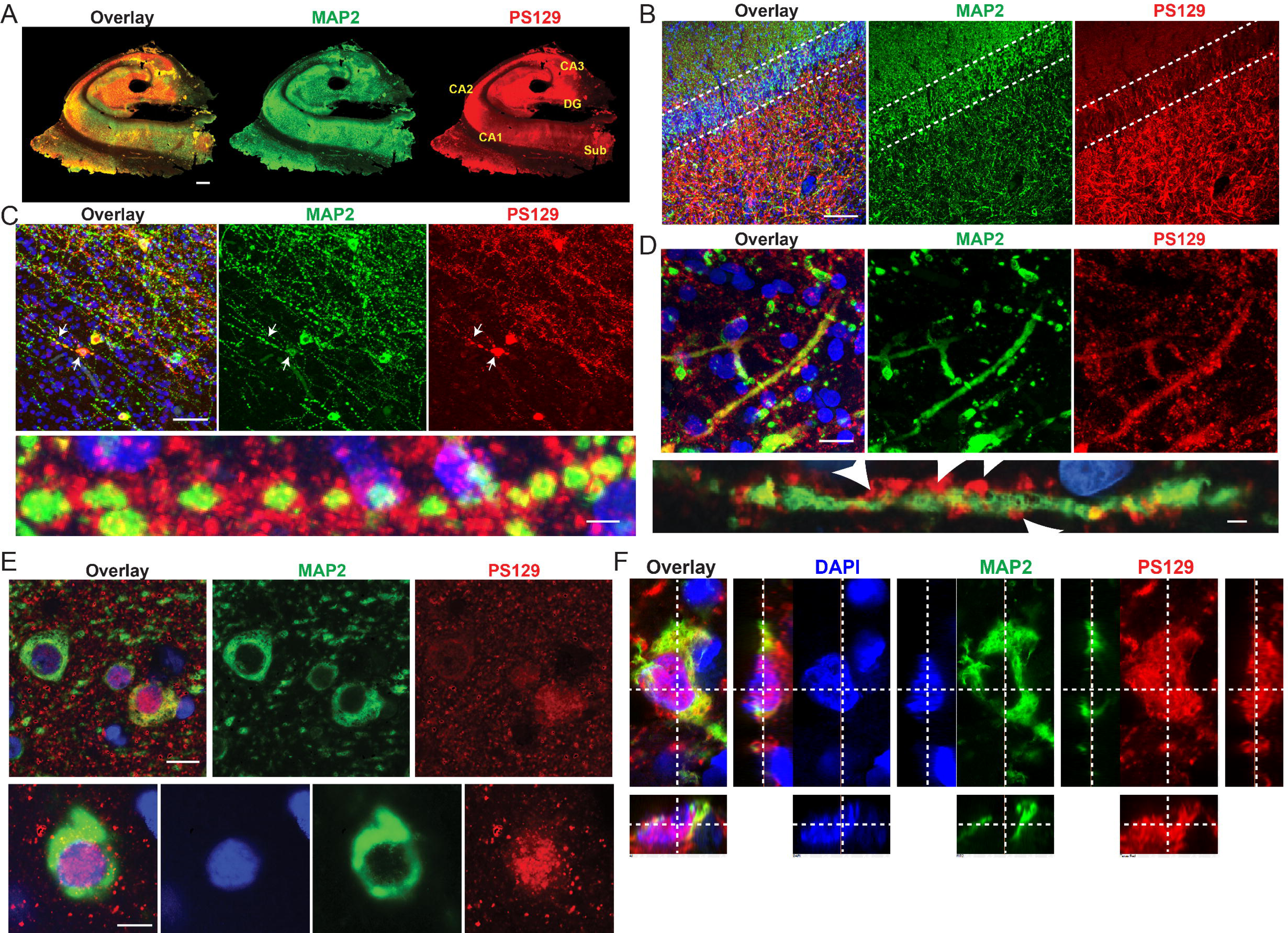
Subcellular localization of PS129 in surgical brain tissue. Surgical TC and HIP sections were immunostained for MAP2 (green), PS129 (red), and DAPI (nuclei). (A) Low-magnification image of the HIP showing predominant colocalization of PS129 with MAP2-positive regions. PS129-enriched hippocampal layers (CA1–3, DG, and Sub) are labeled by name. (B) High-magnification view of the HIP region. Dotted line denoting distinction between hippocampal regions. (C) HIP image showing MAP2-positive varicose dendrites surrounded by PS129-positive puncta. (D) The HIP section displays PS129-positive puncta around normal morphology MAP2-positive dendritic segments. (E). Images take from the TC showing PS129 localization within the nucleus, in the soma, and surrounding the perikarya as punctate structures. (F) Slices’ view of PS129 positive nucleus. Signal distribution for all channels is shown for the y, x, and z planes. Scale bars: A = 1 mm; B,C low mag = 50 µm; C, lower panel 5µm; D upper panel = 20 µm, D lower panel = 2µm; E = 5 µm.

In the hippocampus, varicose dendrites were observed (Fig. 7C), a characteristic “bead on string” appearance, with PS129 between beads and sometimes overlapping MAP2 blebs (Fig. 7C). In several instances, neurons projecting varicose dendrites had perinuclear cytosolic accumulation of PS129 (Fig. 7C, white arrows). High-magnification images confirmed intense PS129 labeling, closely associated with varicose dendrites (Fig. 7C, lower panel). PS129 was also found in close association with dendrites of normal morphology (Figure 7D, continuous MAP2 labeling). PS129 was within these dendrites and appeared decorating the dendritic surface, consistent with dendritic spine localization (Fig. 7D, bottom panel). Some dendrites, varicose or not, lacked PS129 (Fig. 7C-D).

Nuclear PS129 was observed sporadically throughout both the temporal cortex and hippocampus (Fig. 7E-F). MAP2 cells were both positive and negative for nuclear-PS129, and perinuclear PS129 puncta were also commonly observed (Fig. 7E). Orthogonal images through the nucleus of a MAP2-positive cell confirmed that PS129 was inside the nuclear compartment (Figure 7F).

### Human αsyn and PS129 interactomes overlap with cynomolgus macaque

To investigate whether epilepsy influenced the human αsyn and PS129 interactomes, we performed BAR on healthy cynomolgus macaque (CN, Macaca fascicularis) brain tissue that had been pristinely preserved by perfusion fixation. Macaca fascicularis shares 92.83% genome sequence identity with humans(48). CN’s had abundant PS129, particularly in gray matter (Fig. 8A) resembling the human brain (Fig. A, Fig. S2A). Perikarya and neuronal processes, along with abundant diffuse punctate structures, were PS129-positive in the temporal gyrus (Fig. 8A, red box) and the hippocampus (Fig. 8A, blue box). Like human surgical specimens, CIAP pretreatment abolished PS129 reactivity (Fig. 8A, +CIAP).

**Figure 8.**
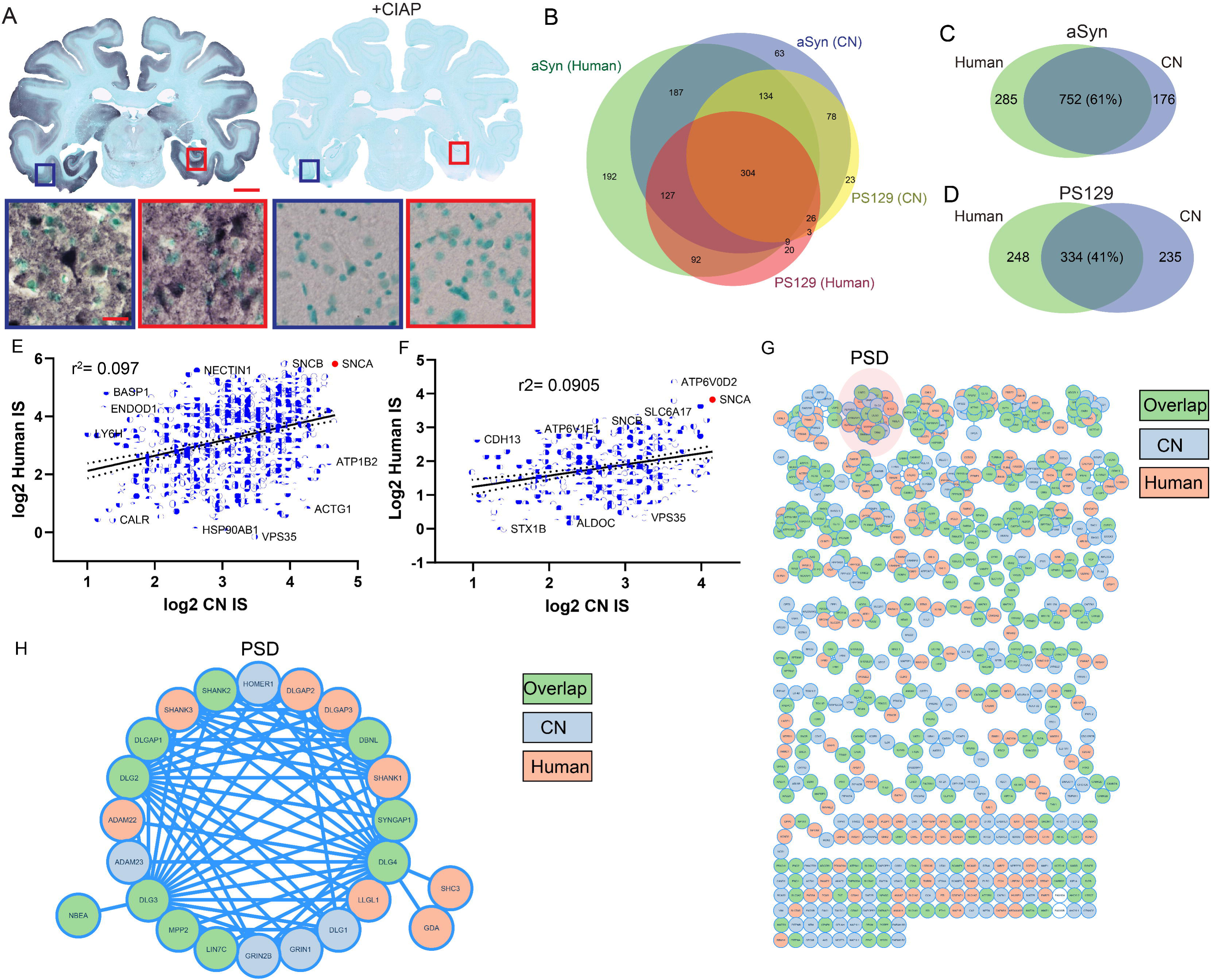
Human and cynomolgus monkey exhibit comparable proteomic patterns. The hippocampus (HIP) and inferior temporal cortex (TC) from a normal cynomolgus monkey (CN, Macaca fascicularis; see Table 2) were characterized for BAR-αSyn and BAR-PS129 protein interactions. A. Coronal brain sections containing TC and HIP were immunostained for PS129 with or without CIAP pre-treatment. Low-magnification images highlight the inferior temporal gyrus (TC; blue box) and HIP (red box); higher-magnification views of the boxed regions are shown. Scale bars: 0.5 mm and 25 μm. B. Tissue lysates from TC and HIP were subjected to BAR-αSyn and BAR-PS129 capture. Proteins identified in the CN were compared with those from human surgical samples using weighted Venn diagrams for both BAR-αSyn and BAR-PS129 interactomes. C. Direct comparison of BAR-αSyn-enriched proteins in human versus CN is displayed in a weighted Venn diagram. The top 10 GO terms for proteins uniquely identified in human (285), cynomolgus monkey (176), or shared between both (752) were determined using g:Profiler and are displayed. (D) A weighted Venn diagram shows the direct comparison of BAR-PS129-enriched proteins in human versus CN. (E) Correlation plots of the 752 overlapping BAR-αSyn proteins and 334 overlapping BAR-PS129 (F) proteins between human and CN are shown; SNCA is highlighted in red. (G) STRING map of physical interactions for BAR-PS129 proteins, color-coded with the data set of origin. The post-synaptic density (PSD) cluster is highlighted. (H) Enlarged PSD cluster shows species-specific proteins within this pathway.

Hippocampus, inferior temporal gyrus, and medial temporal gyri from CN’s were dissected out and subjected to BAR. For CN, BAR-αSyn identified 1,067 proteins, while BAR-PS129 identified 661. Many CN interactions overlapped with human data (Fig. 8B), with 304 proteins shared for all species and BAR-captures. Between species, BAR-αSyn shared 752 proteins (285 human-unique, 315 CN-unique, Fig. 8C), while BAR-PS129 shared 334 proteins (Fig. 8D, 248 human-unique, 327 CN-unique). For proteins found in both species, 448 were unique to αsyn, 30 unique to PS129, and 304 shared (Fig. 8D). For both species, vesicle-related enrichment terms were largely shared, along with glutamatergic synapse, likely reflecting the excitatory neuron–rich composition of the sampled brain regions. 30 PS129 specific proteins were shared between species and were enriched for PSD scaffold proteins (DLG2–4, DLGAP1, SYNGAP1, SYNPO, CASKIN1, GRIA2, IQSEC1/2, CYFIP1/2, SPTBN2, TLN2), presynaptic active zone (BSN, ERC2, PPFIA3, CLTC, MYO5A), nuclear proteins (OGT, CAMKK2, FBXO41), and signaling, secretory, or adhesion regulators (MACF1, DYNC1H1, SPIRE1, WDR7, ADGRB1, AJM1, PKP4, SEC31A). Detailed CN characterization, as well as BAR validation and analyses, are provided in Supplementary Figures 6–7 and Supplementary Table 1.

For BAR-αSyn, we observed moderate correlation between importance scores of humans and CN (r = 0.3122, r^2^=0.09748, p < 0.0001, Fig. 8E). Proteins such as SNCA (highlighted in red), SNCB, NECTIN1, BASP1, ENDOD1, LY6H were more abundant in human than CN. CN. Conversely, in CN, proteins including ATP1B2, ACTG1, VPS35, HSP90AB1, and CALR exhibited higher scores than in humans. For BAR-PS129, we also observed moderate correlation between CN and Human (r=0.3001, r^2^=0.0905, p <0.0001, Fig. 8F). Proteins including ATP6V0D2, SNCA, SV2B, PITPNA, SYT1, and SLC6A17 exhibited noticeably higher scores in humans, whereas VAMP2, WDR7, MYO5A, EEF2, and IQSEC1 displayed higher scores in CN than in humans.

PS129-interactions identified in humans were enriched for postsynaptic processes (Fig. 6), and we next determined whether CN-PS129-interactions were similar to those of humans. To do this, BAR-PS129 identified proteins for both human and CN were plotted in STRING, MCL-clustered (inflation parameter 4), and annotated for whether the protein was found in human, CN, or both (Figure 8G). The large interaction map shows many clusters containing a mixture of proteins identified in human and CN (Figure 8G). All clusters of 4 or more proteins contained a mixture of proteins identified in human and CN, except one cluster containing VAPA, GET3, RAB3GAP2, and HACD3 that was enriched for GO:BP: Protein localization to the endoplasmic reticulum (FDR = 0.0064). PSD proteins were among the top clusters (Fig. 8G, “PSD”) and contained proteins identified in both human and CN. Expansion of the PSD cluster showed a network similar to that identified before (Fig. 6D), with ADAM23, GRIN2B, HOMER1, GRIN1, and DLG1 identified in CN and not in human (Fig. 8H). Together, the interactions identified in CN were similar to those of humans. PS129-specific PSD proteins are present in the healthy primate brain and thus are not specific to disease (epilepsy).

## Discussion

A pivotal technical advancement in this study was the unambiguous detection of physiological, non-aggregated PS129 in surgically resected human temporal lobe tissues. Prior efforts to characterize PS129 were hindered by PMI-driven dephosphorylation, which rapidly removed the phosphoepitope, potentially due to unchecked phosphatase activity in the absence of an ATP-dependent kinase. By leveraging near-zero PMI, via rapid fixation of surgically collected brain specimens, we preserved physiological PS129 at levels in the range of 10-40% of αsyn and demonstrated the conditions in which PS129 can be differentiated from aggregated PS129 (e.g., CIAP pretreatment, Fig. 2). Once we established those conditions, we applied BAR to delineate interactomes of αsyn and PS129 in the intact native human brain tissues. Although presynaptic interactions dominated overall (Fig. 5), unexpectedly, post-synaptic sites and the nucleus had particular significance for PS129 (Fig. 6). These comprehensive proximity interactomes can serve as a “roadmap” and a valuable reference to understand αsyn’s normal functions as well as transition to disease states. Furthermore, the apparent paradox- that the best IHC marker of Lewy pathology is actually a normal, abundant, and highly labile PTM—can now be resolved: pathological PS129 accumulation reflects failure of turnover (dephosphorylation) of a physiological pool rather than (or in addition to) aberrant phosphorylation (21, 23, 24, 29, 31).

### Relevance to **α**syn biology

Physiological αsyn is concentrated at presynaptic nerve terminals(92, 93), where it’s purported to have several functions, including SNARE assembly, vesicle tethering, and regulation of endo/exocytosis(2, 3, 39). Physiological PS129 is abundant in neurons/processes of the hippocampus, olfactory bulb, and substantia nigra (21), largely mirroring αsyn expression(94, 95). PS129 likely acts within processes, driven by neuronal activity, to fine-tune presynaptic transmission(23, 24, 29). Our data support αsyn interactions with synapsin and VAMP2 (7, 24), as well as with other components of the SNARE machinery(5, 96), cytoskeleton(25), which are strengthened with PS129. Other proximity proteomic data have implicated αsyn in mRNA processing and P-body maturation (39, 40); we did not see evidence for these αsyn- or PS129-interactions, but differences between culture and intact human/CN brain, and how αsyn distributes between subcellular compartments, could explain this discrepancy. Within the intact brain, perinuclear and nuclear αsyn was likely overshadowed by synaptic terminal αsyn, making RNA processing functions underrepresented.

PS129 enrichment at post-synaptic densities and the nucleus was unexpected, but not unprecedented. Physiological αsyn has been unequivocally identified in the presynapse but less is known about postsynaptic localization and function(92, 97). Our previous study found physiological PS129 in the apical dendrites of olfactory bulb mitral cells, and PS129 some mitral cell nuclei (21). PS129 interactions with PSD receptors, scaffolding, and kinases (e.g., SHANK1/3, DLGAP1-4, DLG2-4, SYNGAP1, GRIA2/4, IQSEC1/2, etc.), may explain PS129s influence on post-synaptic currents (23). αSyn nuclear localization has been contested(93, 98, 99) but we detected PS129 within nuclei (Fig. 7) as well as PS129-specific enrichment for nuclear proteins (Fig 6, HUWE1, HNRNPM, RBM14, ITCH, OGT, PHF24, and PPP2R5E), therefore human brain nuclei can contain αsyn and PS129. Nuclear localization was confirmed by fluorescent labeling PS129 in human brain (Fig.7). Neuronal nuclei do not contain synaptic vesicles but potentially, αsyn associates with non-synaptic secretory machinery like those involved with neurotrophic signaling (e.g., VGF, SCG2), consistent with generalized αsyn role in endocytic vesicular trafficking(39). PS129 recruitment to DNA damage sites(100–102) or direct αsyn-DNA interactions are possible, but inconsistent with the lack of identified histones interactions.

Kinases that act on S129 include PLK2, PLK3, Casein kinase 1 and 2, GRK2, GRK5, GRK6, LRRK2, GSK-3β(27, 103–108). We identified 118 kinases and 45 phosphatases in human/CN brain close association with αsyn and PS129 (Dataset S3). Of the kinases, GRK2 and GSK-3β were identified, and for phosphatases PPP2R2A was identified. Absence of kinases such as PLK2 suggests they aren’t major contributors to physiological S129 phosphorylation, however they still may have pathological roles, or potentially brain region specificity (e.g., midbrain).

Instead, CAMK2B (PS129’s most abundant kinase) and CASK (i.e., PSD subnetwork, Fig. 5D), both serine kinases with major roles for plasticity at the post-synapse(109, 110).

### Relevance to disease

We confirmed in the human brain that PS129 is a common physiological PTM, not disease-specific (21, 23, 24, 111, 112). Apparent pathological PS129 accumulation results at least in part because dephosphorylation is inhibited by αsyn misfolding/aggregation, thereby selectively preserving PS129 in post-mortem specimens. This interpretation contrasts but does not preclude PS129’s active involvement in synucleinopathy disease processes. When PS129 transitions from enzymatically labile to enzymatically stable will be an essential question to answer.

Our data support the hypothesis that PD mechanisms occur within the confines of normal αsyn biology(37). 57 PD proteins (KEGG:05012, 250 proteins total) were identified for αsyn-interactions, 30 of which were shared with PS129. Therefore, nearly a quarter of PD proteins are canonical αsyn interactors, showing convergence of PD mechanisms on normal αsyn biology. Strikingly, PS129-unique PD proteins were not identified, highlighting a possible disconnect between physiological and pathological-PS129 interactomes. Proteins identified from known monogenetic PD genes (SNCA, VPS35, DJ-1, DNJC6, and SYNJ1) were mostly shared between αsyn and PS129, except for DJ-1, which was unique to αsyn.

Previous BAR-PS129 in the synucleinopathy brain identified presynaptic vesicle processes (35), similar to here, suggesting pathological aggregation occurs within the context and physical space of normal αsyn biology. Pre- and postsynaptic αsyn aggregation may result in impaired or loss-of-function, impacting synaptic plasticity, SNAREs, neurotransmitter release, and vesicle trafficking(113–115). Early aggregation has been observed in postsynaptic neurons and presynaptic nerve terminals(116), and therefore misfolding at those sites would kinetically “trap” PS129 in its physiological state, ultimately leading to its accumulation in Lewy bodies.

αSyns role for epilepsy is unclear despite overlapping mechanisms between neurodegenerative diseases (e.g., PD) and epilepsy(117). Intriguingly, the PS129-interactome was associated with several key epilepsy factors (Fig. 6, e.g., DLG4, CASK, GRIA2)(118) showing that physiological-PS129 acts within epilepsy pathways. However, many of these interactions were identified in healthy CN and are thus not specific to epilepsy. PS129 was associated with varicose dendrites (Fig. 7), a pathological feature of epileptic seizures/excitotoxicity(119), as well as dendrites of normal morphology similar to what we previously observed in healthy mice(21).

For a few neurons with varicose dendrites, PS129 accumulation was apparent in the perikarya. We found no evidence that αsyn was aggregated (i.e., enzymatically resistant) in epilepsy tissues (Fig. 2). Interestingly, PLK2, which can phosphorylate αsyn(103, 104) but was not identified here, mediates the degradation of postsynaptic proteins, including PSD95, as a neuroprotective response to reduce AMPA receptor (identified here) expression in epileptic patients. Together, our data support αsyn/PS129 acting within epilepsy pathways, potentially the downregulation of post-synaptic AMPA receptors(120), but the mechanisms are likely distinct from the aggregation-driven neuropathology of synucleinopathies.

### Limitations

PS129 abundance and distribution are driven by neuronal activity (23), which could have been altered in intractable epilepsy, as observed for tau (121, 122). However, several of our findings refute this interpretation, including 1) human PS129 interactomes overlapped well with CN, 2) and PS129 abundance/distribution in surgical tissues similar to CN and mice.

Nevertheless, we cannot completely exclude the possibility that epilepsy impacted αsyn interactomes. But we can confidently assign interactomes to non-aggregated αsyn, and, outside the context of pathological aggregation, this is an impossible feat without surgically resected tissues. Another limitation was the cross-species analysis, which was restricted to proteins with high interspecies homology, potentially underestimating interspecies interactome overlap. A technical limitation, PS129-specific interactions may reflect rare PS129 interactions, due to biased peptide selection during DDA, that would “boost’ less prominent interactions that were otherwise excluded. Lastly, multiple specimens from one CN monkey with prominent PS129 were used to benchmark human interactions, sufficient to identify many significant interactors (1067 for αsyn and 661 for PS129), but likely insufficient to map the CN interactome comprehensively. Lower mammalian interactomes have been detailed (21, 39, 123), and the focus of this study was to determine the human interactome.

### Future Directions

As with all promiscuous labeling proximity proteomic methods, we can only infer interactions, but we cannot, and do not make any claims about direct protein-protein interactions, which must be investigated with follow-up studies. With completion of this study, we now have physiological interactomes and pathological interactomes in the human brain(34, 35), the next step will be to determine differential interactions between the two states, and how those interactions evolve during pathological αsyn aggregation.

### Concluding remarks

Our results indicated that PS129 is a common PTM in the human brain with an unappreciated significance in postsynaptic neurons. αsyn/PS129 interactomes were highly interconnected with clear biological relevance, but far too large to exhaustively analyze on a protein level basis. Although, αsyn/PS129 presynaptic vesicles interactions are dominant, considerations of non-synaptic endocytic vesicular processes (e.g., VGF signaling) will help unify αsyn interactomes across diverse cell types, contexts, and subcellular compartments of the brain.

## Data availability

### Raw MS data at ProteomeXchange [accession]; TBD Acknowledgements

We want to thank our donors, their families, and the brain banks for providing us with valuable tissue samples. Thank you to the Rush Movement Disorders Brain Bank and Banner Sun Health Research Institute. Funding was provided by NINDS and NIA 5R01NS128467

## Author Contributions

S.G.C. performed studies, analyzed data, prepared figures, and wrote the manuscript. J.B.D., A.B., and T.T. performed experiments and acquired data. S.S. collected surgical tissues. J.H.K. helped prepare the manuscript. S.M. performed studies. B.A.K. performed studies, conducted data analysis, constructed figures, and wrote the manuscript.

