## Supplementary material for "Physiological α-synuclein S129 phosphorylation mediates postsynaptic and nuclear interactions in the human brain": SI

PNAS

**This PDF file includes:**

**Figure S1**. Physiological PS129 is undetectable in FFPE human post-mortem hippocampus with short post-mortem intervals (PMI 2–3 hours).
**Figure S2**. PS129 IHC in surgical and PD/DLB brain tissues.
**Figure S3**. Physiological PS129 is not detectable in FFPE sections of surgical human brain tissues.
**Figure S4**. Protein analysis of PS129 in surgical and PD/DLB brain specimens.
**Figure S5**. Merged BAR-αSyn and BAR-PS129 physical interaction map.
**Figure S6**. BAR-αSyn and BAR-PS129 interactomes in non-human primate brain
**Figure S7**. Searching CN peptides against only the Macaca fascicularis UniProt FASTA database yielded a limited number of protein identifications.
**Figure S8**. c-Fos IHC in surgical human and CN hippocampal and cortical tissues.
**Table S1**. Combined wet mass of surgical and CN samples used for BAR captures.

**Other supporting materials for this manuscript include the following:**

**Dataset S1 (separate file).** Surgical samples – full proteomics dataset (LFQ Analyst).
**Dataset S2 (separate file).** BAR-αSyn- and BAR-PS129-enriched proteins ranked by importance.
**Dataset S3 (separate file).** Phosphatases and kinases identified in surgical and CN proteomics datasets.
**Dataset S4 (separate file).** CN samples – full proteomics dataset (LFQ Analyst) and BAR-enriched proteins ranked by importance.

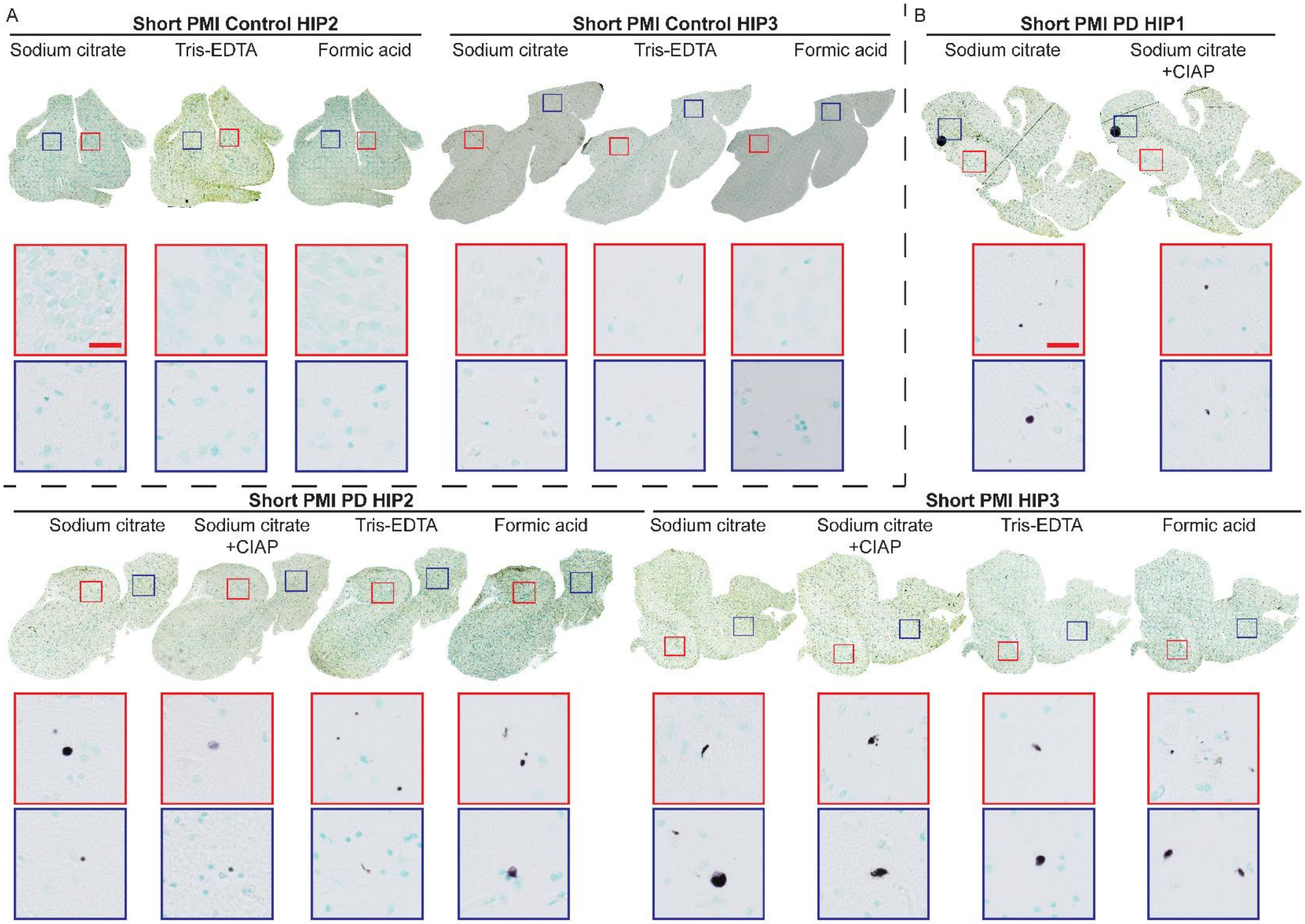


**Fig. S1.** Physiological PS129 is undetectable in FFPE human post-mortem hippocampus with short post-mortem intervals (PMI 2–3 hours). IHC staining for PS129 was performed in hippocampal sections from **A** control (non-synucleinopathy) cases and **B** PD cases with markedly short PMIs. Tissues underwent antigen retrieval using sodium citrate, Tris-EDTA, or formic acid prior to PS129 staining. Sections underwent antigen retrieval with sodium citrate, Tris-EDTA, or formic acid prior to PS129 staining. For PD tissues, CIAP pretreatment was applied before antigen retrieval to distinguish pathological from physiological PS129. No PS129 immunoreactivity was observed in control hippocampus under any retrieval condition. In PD cases, PS129-positive structures remained resistant to CIAP treatment. Scale bars= 25 µm (applies to all high-magnification panels).

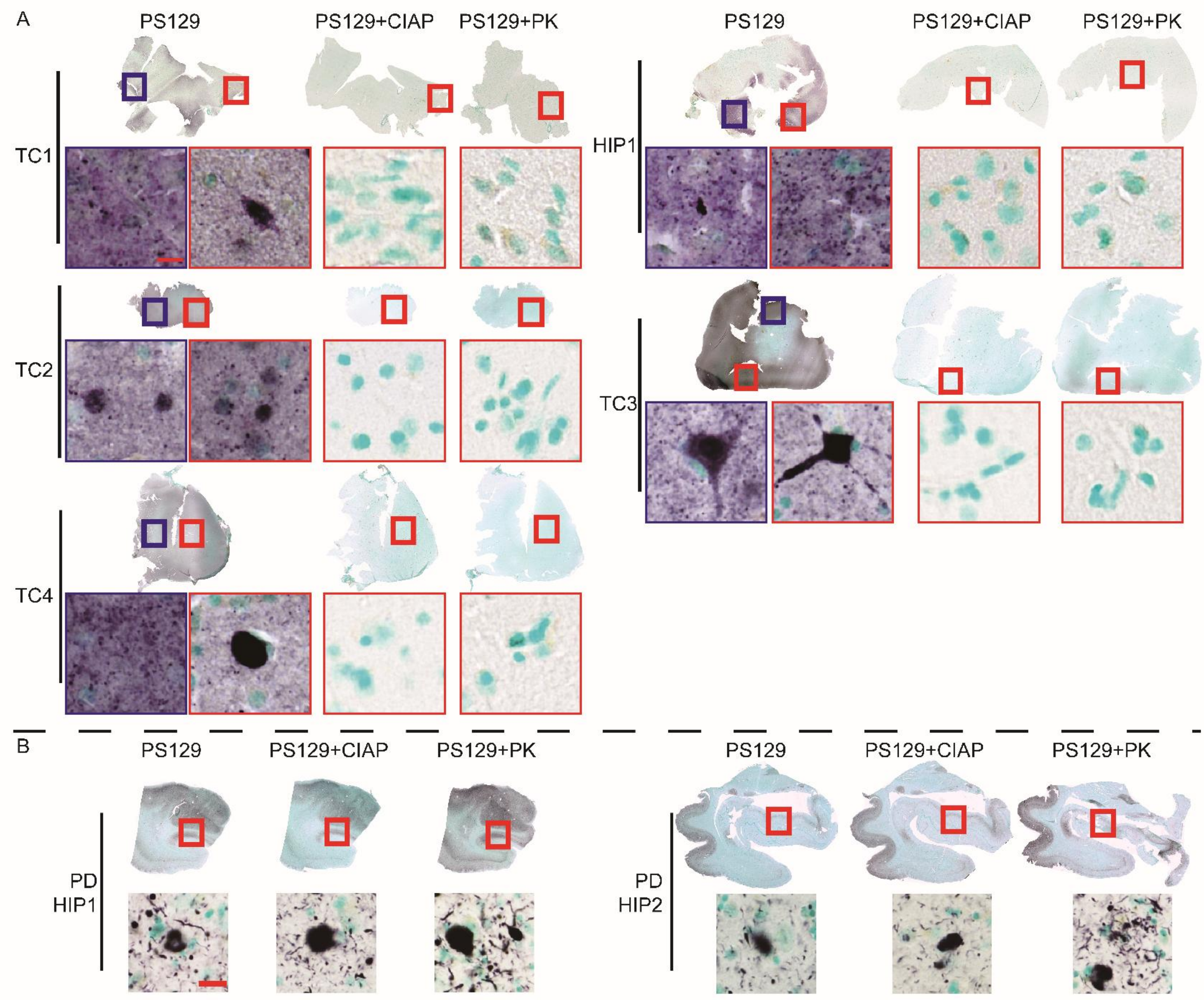


**Fig. S2.** PS129 IHC in surgical and PD/DLB brain tissues. Whole-section scans and representative higher-magnification (enlarged 20×) images of **A** surgical temporal cortex and hippocampus and **B** post-mortem PD/DLB hippocampus immunostained for PS129, with or without prior CIAP or PK treatment. Boxes (red or blue) in whole-section images indicate the regions shown at higher magnification. In surgical specimens, PS129 immunoreactivity was completely abolished by both CIAP and PK pretreatment. In contrast, PD/DLB tissues exhibited CIAP- and PK-resistant PS129 staining; PK pretreatment enhanced PS129 signal intensity. Scale bars= 10 μm (applies to all high-magnification panels).

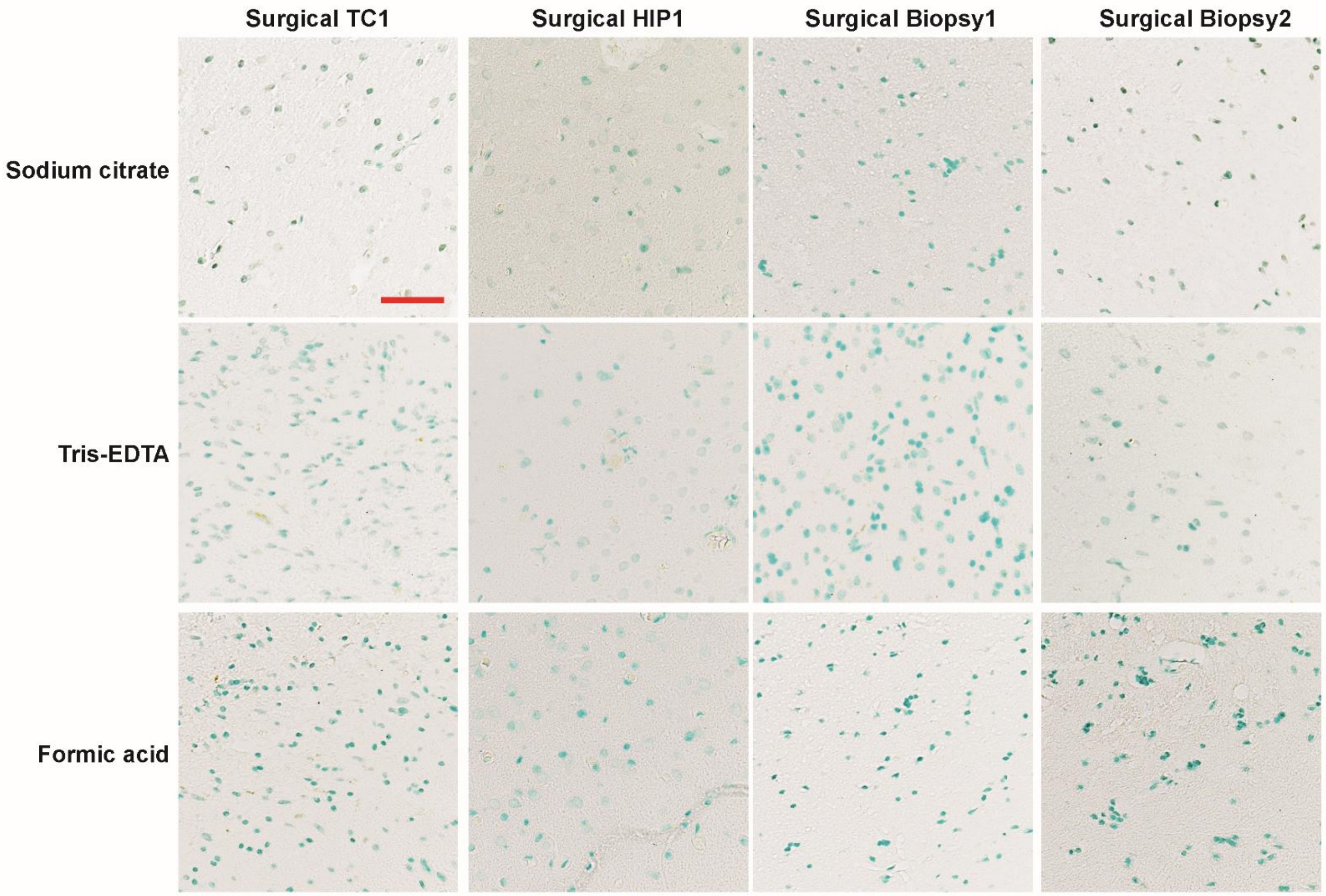


**Fig. S3.** Physiological PS129 is not detectable in FFPE sections of surgical human brain tissues. Immunohistochemical staining for PS129 in surgically resected temporal cortex and hippocampus, as well as biopsies of temporal cortex, from the same patient (Patient ID: 1, Table 1). All tissue was fixed in 4% PFA within 30 minutes of resection. Large surgical specimens were cryoprotected in 30% sucrose, frozen-sectioned, and the remaining tissues subsequently processed for paraffin embedding after rinsing and storage in 70% ethanol (Surgical TC1 and HIP1). Small biopsies (TC Biopsy 1 and 2) were directly paraffin-embedded after PFA fixation. All FFPE sections were cut at 4 µm thickness and subjected to antigen retrieval using sodium citrate, Tris-EDTA, or formic acid prior to PS129 immunostaining. No PS129 immunoreactivity was observed under any condition. Scale bar: 50 µm (applies to all panels).

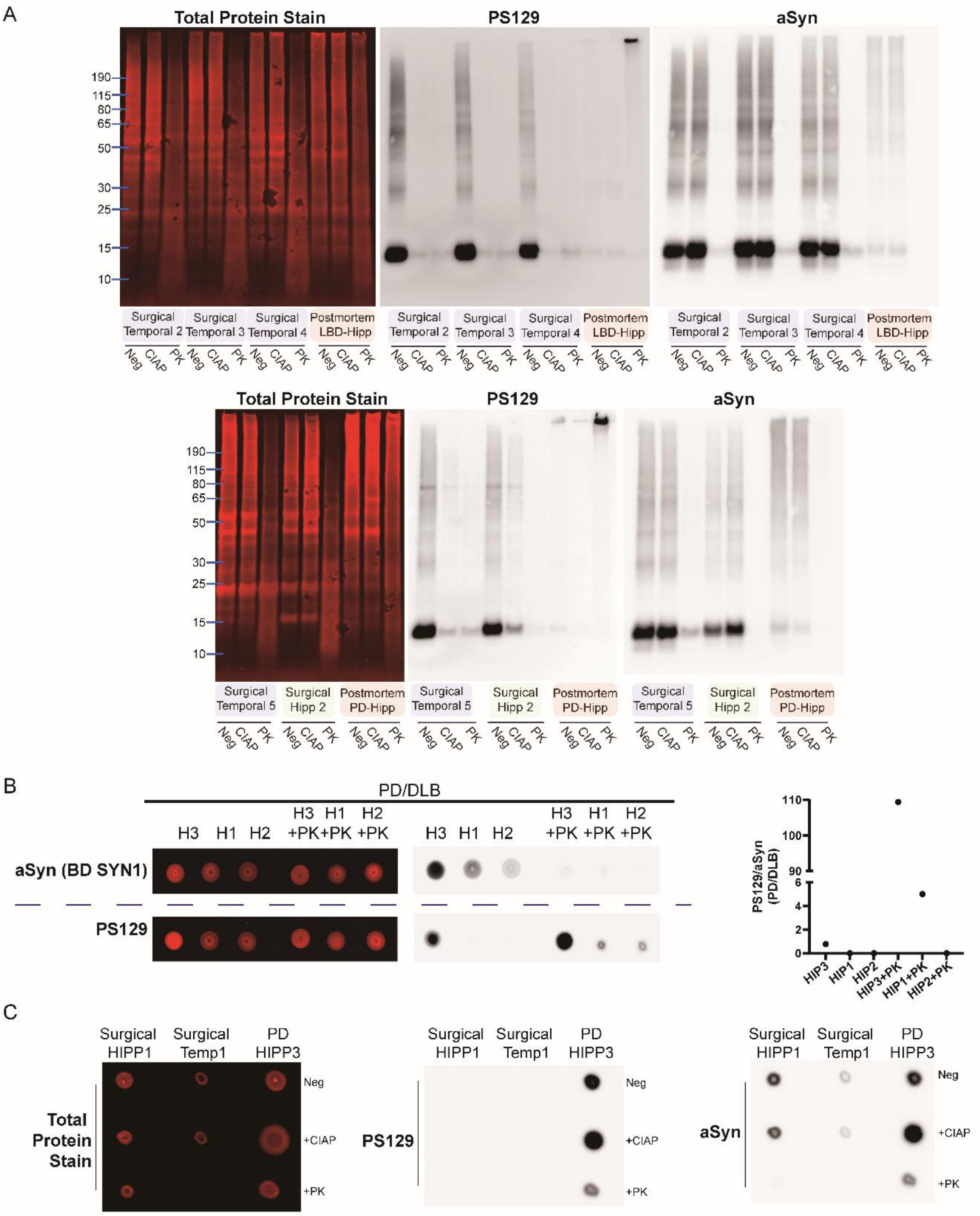


**Fig. S4.** Protein analysis of PS129 in surgical and PD/DLB brain specimens. **A** Tissue lysates (10 µg) from surgical and PD/DLB cases were separated on 4–12% Bis-Tris gels, transferred to PVDF membranes, and probed for total protein, PS129, or αSyn (BD SYN1). WB results correspond to the cases shown in Figure 1. Neg, no enzymatic pretreatment; CIAP, CIAP pretreatment; PK, PK pretreatment. PS129 blot images were adjusted for brightness/contrast to visualize weak signals. **B** The PS129:αSyn (BD SYN1) ratio in PD/DLB hippocampal lysates was quantified by dot blot and is presented as a scatter plot. **C** Insoluble pellets remaining from surgical HIP1, TC1, and PD HIP3 samples after initial lysis were solubilized in 8 M urea buffer and analyzed by dot blot for PS129 and αSyn (BD SYN1).

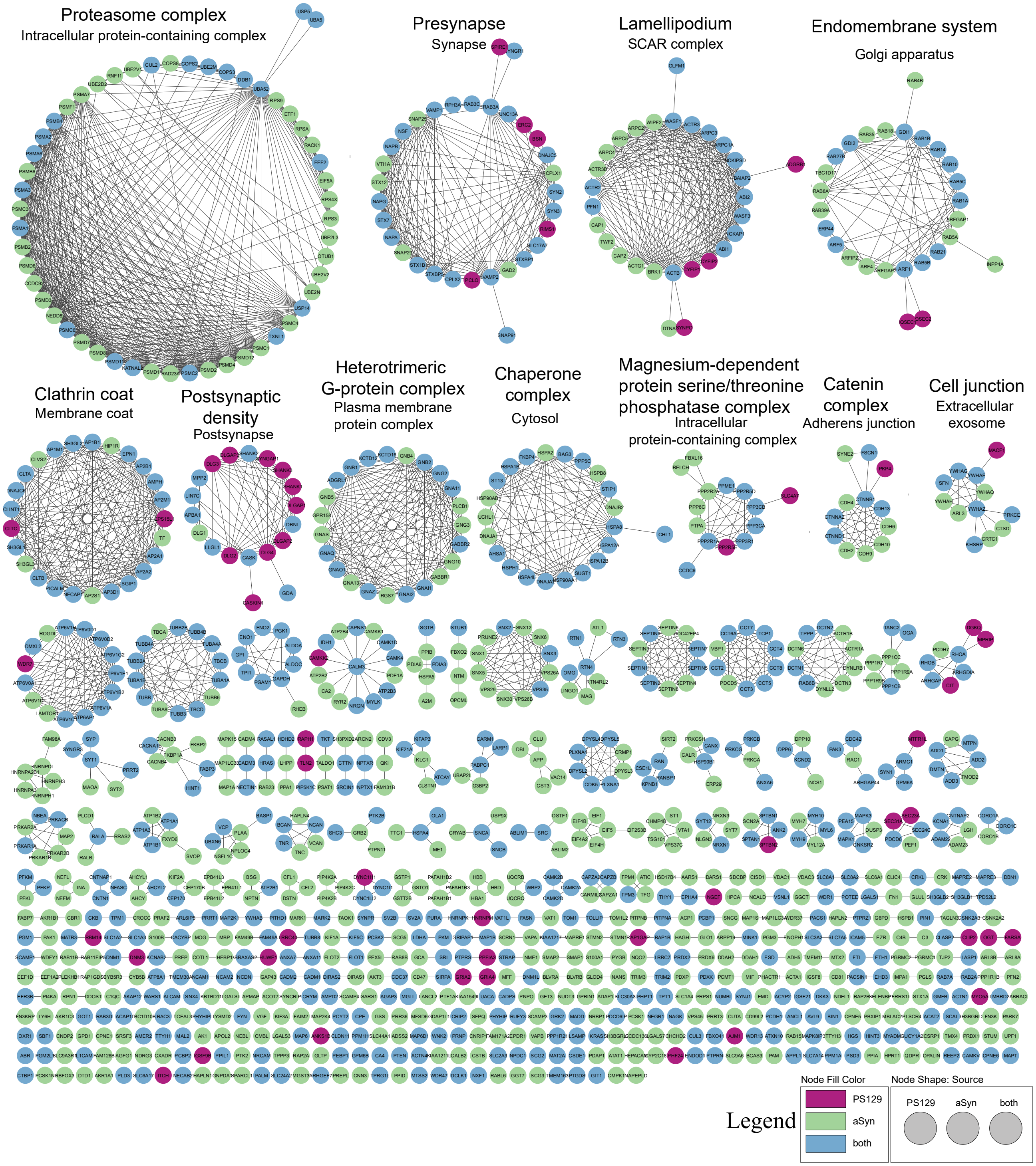


**Fig. S5**. Merged BAR-αSyn and BAR-PS129 physical interaction map. Proteins significantly enriched in BAR-αSyn and BAR-PS129 relative to background controls (BAR-Neg; adjusted $p < 0.05$) were analyzed. The lists of BAR-αSyn– and BAR-PS129–enriched proteins were imported into Cytoscape, and physical interaction subnetworks for each condition were generated using STRING (confidence score ≥ 0.4; FDR ≤ 5%). The two resulting protein–protein interaction (PPI) networks were merged and subjected to Markov Cluster (MCL) algorithm clustering. Node colors indicate BAR-αSyn–specific (green), BAR-PS129–specific (pink), or shared (blue) proteins. The largest clusters were manually annotated with their most significant GO Cellular Component (GO:CC) terms; font size reflects statistical significance ($-\log_{10}$ adjusted p-value). Edge thickness corresponds to the STRING confidence score.

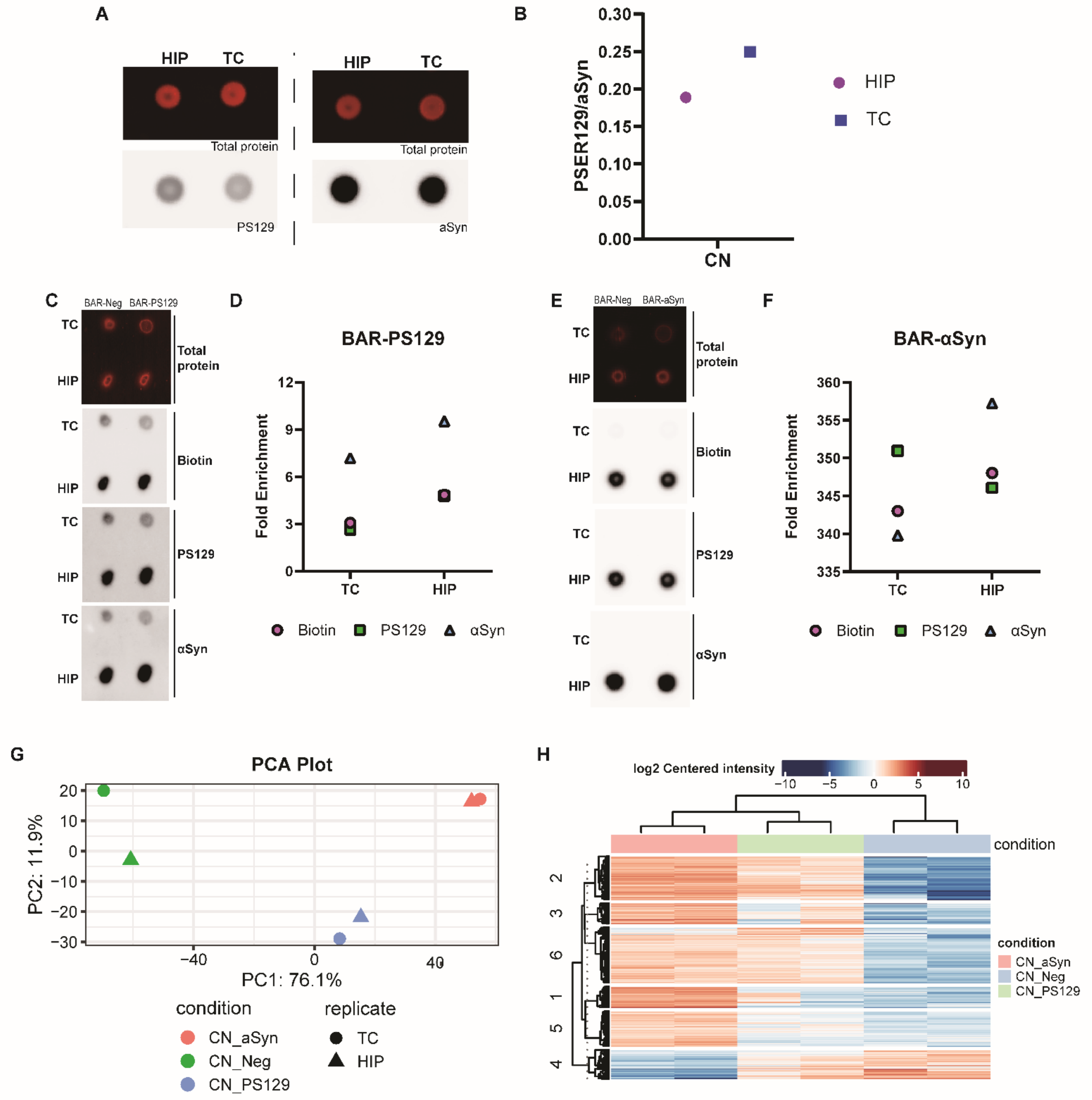


**Fig. S6.** BAR-αSyn and BAR-PS129 interactomes in non-human primate brain. Hippocampus (HIP) and inferior/medial temporal gyrus (TC) from a normal cynomolgus monkey (Macaca fascicularis) were analyzed. **A** Lysates (1 μL) from HIP and TC were spotted onto PVDF membranes and probed for total protein, PS129, and αSyn (BD SYN1). Signals were acquired under optimal exposure using a ChemiDoc imager and quantified with ImageJ. **B** PS129:αSyn ratios are shown as scatter plots. **C-F** HIP and TC lysates were subjected to BAR-PS129 and BAR-αSyn capture. Bead eluents, together with BAR-Neg controls, were spotted onto membranes and probed for total protein, biotin, PS129, and αSyn. Signals were quantified with ImageJ, normalized to the mean of BAR-Neg controls, and plotted. Eluents were further analyzed by LC-MS/MS. **G** PCA plot and **H** hierarchical clustering heatmap of CN BAR-Neg, BAR-αSyn, and BAR-PS129 samples are shown.

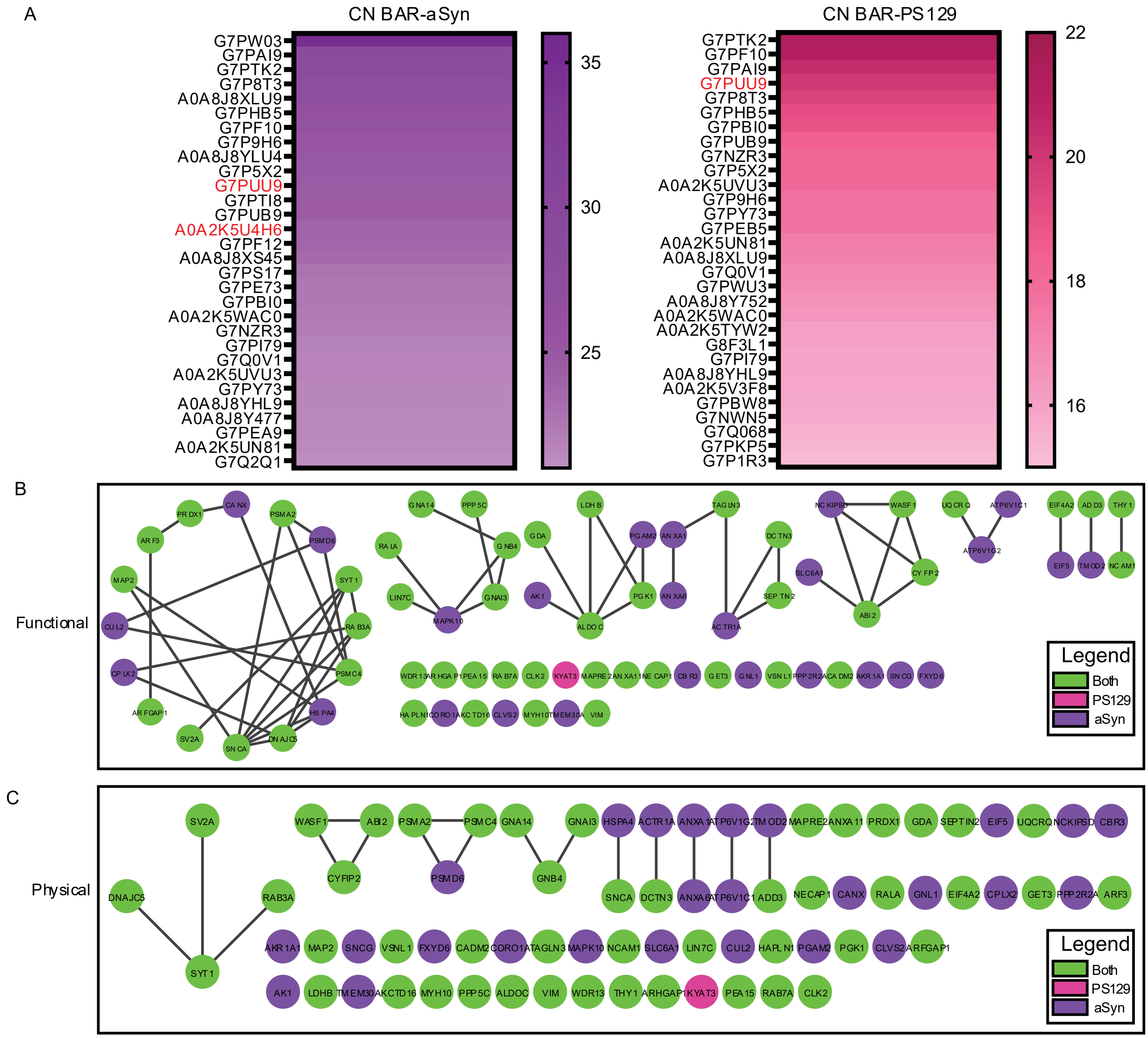


**Fig. S7.** Searching CN peptides against only the *Macaca fascicularis* UniProt FASTA database yielded a limited number of protein identifications. Raw files from the CN BAR LC-MS/MS run were searched against the *Macaca fascicularis* proteome (UniProt). Using LFQ Analyst ($\log_2$ fold-change cutoff ≥ 0, adjusted $p \leq 0.05$), 754 proteins were quantified. However, the majority of these entries were unreviewed UniProt/TrEMBL proteins scheduled for removal in early 2026 because they are not part of the reference proteome (e.g., entry G7P1R3 is flagged: “Entry scheduled for removal from UniProtKB/TrEMBL as it is not part of a reference proteome”). **A** Among the top 30 enriched proteins in the BAR-αSyn and BAR-PS129, only two (G7PUU9 and A0A2K5U4H6, highlighted in red) were reviewed entries; all others were unreviewed. **B** Consequently, when the full protein lists were imported into Cytoscape and mapped to functional or physical interactions via STRING, the resulting networks contained very limited number of identified proteins in either condition, rendering the dataset unsuitable for downstream analysis.

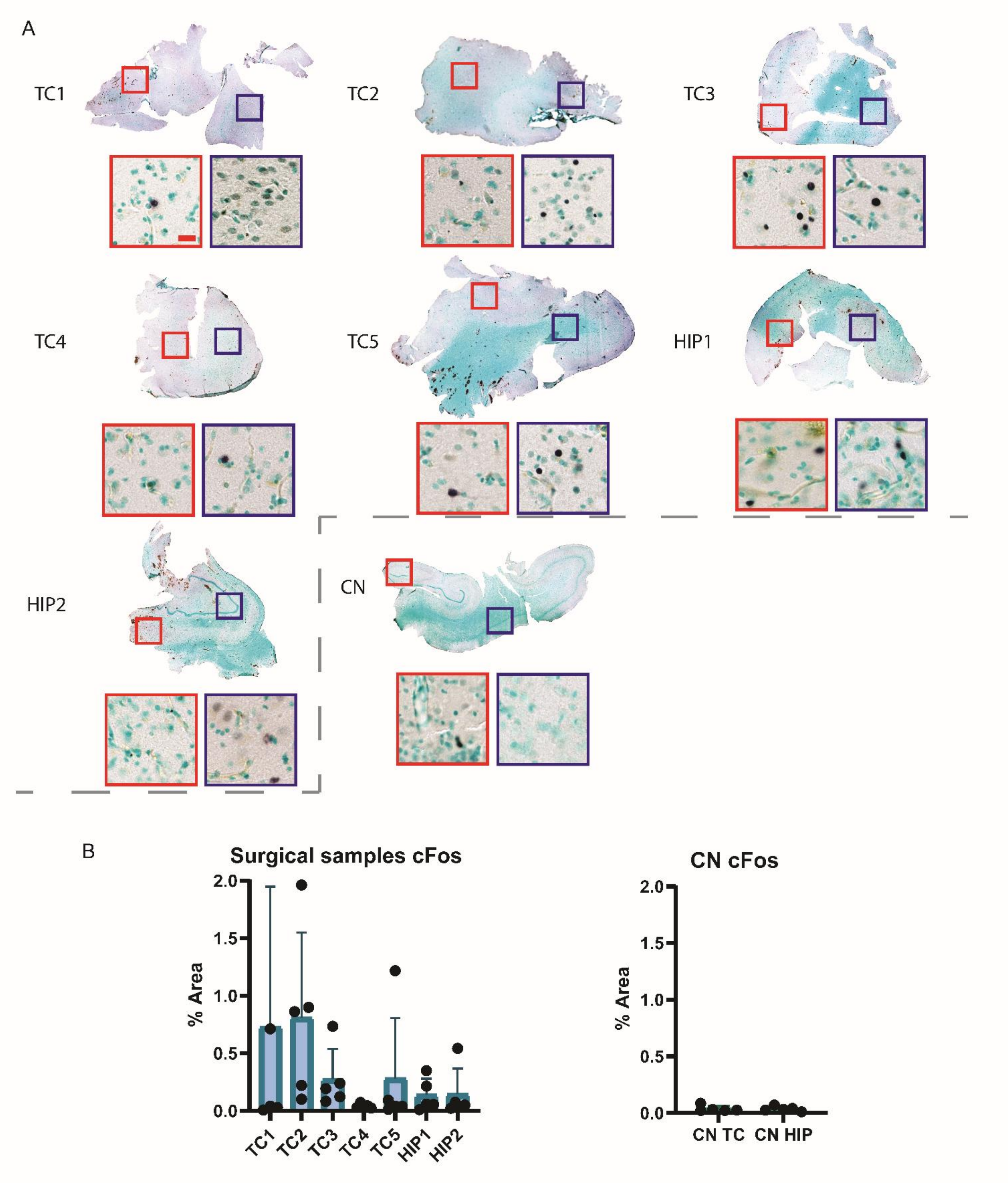


**Fig. S8.** c-Fos IHC in surgical human and CN hippocampal and cortical tissues. Whole-section scans and corresponding higher-magnification (enlarged 20×) images (regions indicated by red and blue boxes) are shown for **A** surgical human cases and CN hippocampus and temporal gyrus. Scale bar: 20 µm (applies to all high-magnification panels). **B** Five non-overlapping regions were randomly selected from each tissue section, imaged, and c-Fos-positive nuclei were quantified using Ilastik pixel classification and ImageJ analysis. Quantified data are presented in the accompanying graph.

**Tables**

**Table S1.** Combined wet mass of surgical and CN samples used for BAR captures.

| Surgical | | | | | | | |
|---|---|---|---|---|---|---|---|
| **Wet mass (g)** | TC1 | TC2 | TC3 | TC4 | TC5 | HIP1 | HIP2 |
| NC | 0.0203 | 0.0127 | 0.0287 | 0.0162 | 0.014 | 0.0276 | 0.0135 |
| PS129 | 0.0213 | 0.0103 | 0.0257 | 0.0179 | 0.0179 | 0.0251 | 0.011 |
| aSyn | 0.0191 | 0.0079 | 0.0191 | 0.0139 | 0.0181 | 0.0261 | 0.0097 |

| | |
|---|---|
| **Average** | **0.017909524** |
| **STDEV** | **0.006180607** |

| CN | | |
|---|---|---|
| Wet mass (g) | CN TC | CN HIP |
| NC | 0.012 | 0.0121 |
| PS129 | 0.0138 | 0.0116 |
| aSyn | 0.0092 | 0.0114 |

| | |
|---|---|
| **Average** | **0.011683** |
| **STDEV** | **0.001484** |

**Dataset S1 (separate file):** Full proteomics dataset from surgical human hippocampus and temporal cortex samples (downloaded from LFQ Analyst). This file contains the complete LFQ dataset and the protein lists used for the Venn diagrams presented in the main figures.

**Dataset S2 (separate file):** All proteins enriched in surgical BAR-αSyn or BAR-PS129 are ranked by an importance score calculated as: absolute $\log_2$ fold-change × $-\log_{10}$(adjusted *p*-value).

**Dataset S3 (separate file):** Kinases and phosphatases identified in both surgical and CN proteomics datasets. All kinases and phosphatases detected in the surgical and CN datasets are listed. These were subsequently filtered to include only serine/threonine-specific kinases and phosphatases, along with a few closely associated proteins.

**Dataset S4 (separate file):** Full proteomics dataset of CN hippocampus and temporal cortex samples (downloaded from LFQ Analyst). This complete dataset contains peptides identified against both *Homo sapiens* and *Macaca fascicularis* databases. CN BAR-αSyn– and BAR-PS129–enriched proteins are ranked by an importance score calculated as absolute $\log_2$ fold-change × $-\log_{10}$(adjusted *p*-value).
