## Supplementary material for "Physiological α-synuclein S129 phosphorylation mediates postsynaptic and nuclear interactions in the human brain": table 1

**Table 1**. Table of human specimens.

| Patient ID | Tissue ID | Age | Sex | Primary Diagnosis | *H&Y Scale | Disease Duration | Procedure | Brain Region | Fixation protocol | Tissue Sectioning |
|---|---|---|---|---|---|---|---|---|---|---|
| 1 | TC1 | 30 | F | Localization-related (focal) (partial) idiopathic epilepsy and epileptic syndromes with seizures of localized onset, not intractable, without status epilepticus (CMS-HCC) | NA | Perinatally | Left craniotomy temporal lobectomy with ECOG | Tempor al cortex | Immediate immersion fixation in 4% PFA for over 72 hours | 40-micron free-floating/ 4-micron on-slide paraffin-embedded |
| | HIP1 | | | | | | | Hippoca mpus | | |
| | TC Biopsy 1 | | | | | | | Tempor al cortex | Immersed in saline (<30 min) followed by fixation in 4% PFA for over 72 hours. | 4-micron on-slide paraffin-embedded |
| | TC Biopsy 2 | | | | | | | | | |
| 2 | TC2 | 31 | F | Localization-related (focal) (partial) idiopathic epilepsy and epileptic syndromes with seizures of localized onset, not intractable, without status epilepticus (CMS-HCC) | NA | Age 24 onset. Daily Seizures | Left Temporal Lobectomy, Amygdalohippocamp ectomy | Tempor al cortex | Immersed in saline (<3 min) followed by fixation in 4% PFA for over 72 hours. | 40-micron free-floating |
| 3 | TC3 | 39 | M | Partial epilepsy, with intractable epilepsy, pharmacoresistant (CMS-HCC) | NA | Age 16 onset. Monthly | NA | Tempor al cortex | Immersed in saline (<3 min) followed by fixation in 4% PFA for over 72 hours. | 40-micron free-floating |
| 4 | TC4 | 36 | M | NA | NA | Age 19 onset. Seizures several times a month (2-3) | Left craniotomy with temporal lobectomy, hippocampectomy, removal of the left hippocampus electrode, preservation of the left insula RNS electrode, and ECOG. | Tempor al cortex | Immersed in saline (<3 min) followed by fixation in 4% PFA for over 72 hours. | 40-micron free-floating |
| | HIP2 | | | | | | | Hippoca mpus | | |
| 5 | TC5 | 43 | M | Intractable generalized epilepsy (CMS-HCC) | NA | Since 2013 | Right craniotomy temporal lobectomy | Tempor al cortex | Immersed in saline (<15 min) | 40-micron free-floating |

| | | | | | | | | | | |
|---|---|---|---|---|---|---|---|---|---|---|
| | | | | | | | with amygdalohippocampectomy | | followed by fixation in 4% PFA for over 72 hours. | |
| 6 | PD HIP1 | 84 | M | PD | 4 | [+]4 years | NA | Hippocampus | The whole brain immersion fixation in 4% PFA for 7–10 days. PMI: >6 hours | 40-micron free-floating |
| 7 | PD HIP2 | 73 | M | LBD | 2 | [+]11 years | NA | Hippocampus | The whole brain immersion fixation in 4% PFA for 7–10 days. PMI: >6 hours | 40-micron free-floating |
| 8 | PD HIP3 | 81 | F | PD s/p STN DBS | 5 | [+]10 years | NA | Hippocampus | The whole brain immersion fixation in 4% PFA for 7–10 days. PMI: >6 hours | 40-micron free-floating |
| 9 | Short PMI Control HIP1 | ≥90 | F | NA | NA | NA | NA | Hippocampus | The whole brain immersion fixation in 4% PFA. PMI=2.08 hours | 6-micron on-slide paraffin-embedded |
| 10 | Short PMI Control HIP2 | 88 | M | NA | NA | NA | NA | Hippocampus | The whole brain immersion fixation in 4% PFA. PMI=2.63 hours | 6-micron on-slide paraffin-embedded |
| 11 | Short PMI Control HIP3 | 86 | M | NA | NA | NA | NA | Hippocampus | The whole brain immersion fixation in 4% PFA. PMI=2.38 hours | 6-micron on-slide paraffin-embedded |
| 12 | Short PMI PD HIP1 | 83 | F | PD | NA | 5 | NA | Hippocampus | The whole brain immersion fixation in 4% PFA. PMI=2.45 hours | 6-micron on-slide paraffin-embedded |

| | | | | | | | | | | |
|---|---|---|---|---|---|---|---|---|---|---|
| 13 | Short PMI PD HIP2 | 83 | M | PD | NA | 21 | NA | Hippocampus | The whole brain immersion fixation in 4% PFA. PMI=2.47 hours | 6-micron on-slide paraffin-embedded |
| 14 | Short PMI PD HIP3 | 72 | M | PD | NA | 12 | NA | Hippocampus | The whole brain immersion fixation in 4% PFA. PMI=2.65 hours | 6-micron on-slide paraffin-embedded |

*Hoehn and Yahr Scale. +Based on clinical records from the Rush Movement Disorder program, the disease duration measures the time period between initial symptom observation/detection and death.
