## Supplementary material for "Physiological α-synuclein S129 phosphorylation mediates postsynaptic and nuclear interactions in the human brain": table 2

**Table 2**. Table of animal specimens.

| ID | Species | Age at death | Sex | Treatment | Fixation protocol | Tissue Sectioning |
|---|---|---|---|---|---|---|
| Monkey 1 | Macaca fascicularis | 4 years | M | None | Within approximately 5 minutes of anesthesia/euthanasia, animals were transcardially perfused with 4% PFA. | 40-μm free-floating |
| Mouse 1 | C57BL/6J | 4-5 months | M | None | Transcardial perfusion with PBS until the perfusate exiting a small incision in the right atrium was clear, followed by fixation in 4% PFA. | Half: 40-μm free-floating; Half: 4-μm on-slide paraffin-embedded |
| Mouse 2 | C57BL/6J | 4-5 months | F | None | Transcardial perfusion with PBS until the perfusate exiting a small incision in the right atrium was clear, followed by fixation in 4% PFA. | Half: 40-μm free-floating; Half: 4-μm on-slide paraffin-embedded |
| Mouse 3 | C57BL/6J | 4-5 months | F | None | Transcardial perfusion with PBS until the perfusate exiting a small incision in the right atrium was clear, followed by fixation in 4% PFA. | Half: 40-μm free-floating; Half: 4-μm on-slide paraffin-embedded |
